# Going around the bend to understand the role of leg coalescence in metachronal swimming

**DOI:** 10.1101/2024.07.24.605009

**Authors:** Nils B. Tack, Sara O. Santos, Monica M. Wilhelmus

## Abstract

Many of the most abundant aquatic invertebrates display metachronal swimming by sequentially beating closely spaced flexible appendages. Common biophysical mechanisms like appendage spatial asymmetry and phase drive the success and performance of this locomotor mode, which is generally explained by the need to maximize thrust production. However, the potential role of these mechanisms in drag reduction, another important contributor to the overall swimming performance, has yet to be evaluated. We present a comprehensive overview of the morphological, functional, and physical mechanisms promoting drag reduction during metachronal swimming by exploring appendage differential bending and leg grouping (coalescence). We performed μ-CT and *in-vivo* velocimetry measurements of shrimp (*Palaemonetes vulgaris*) to design a five-legged robotic metachronal analog. This test platform enabled simultaneous flow and force measurements to quantify the thrust and drag forces produced by flexible and stiff pleopods (legs) beating independently or coalescing. We tested the hypothesis that coalescence and bending effectively reduce drag during the recovery stroke (RS). The curved cross-section of the pleopods enables passive asymmetrical bending during the RS to reduce their coefficient of drag by up to 75.8% relative to stiff pleopods. Bending promotes physical interactions facilitating the coalescence of three pleopods at any time during the RS to reduce drag such that the mean net thrust produced during coalescence is increased by 30.2%. These improvements are explained by the production of a weaker wake compared with stiff and non-coalescing pleopods. Our results describe fundamental biological and physical components of metachronal propulsion that may aid the development of novel bio-inspired underwater vehicles.

**Summary statement:** Shrimp swimming legs bend nearly horizontally and cluster together during metachronal propulsion to reduce drag and improve the overall swimming performance.

## INTRODUCTION

Metachronal propulsion consists of sequentially beating closely spaced appendages to generate thrust. It is employed by a formidable number of aquatic species across a wide scale spectrum, from microscopic protozoa to large lobsters. Its ubiquity across taxa, including many of the major groups performing mass diel vertical migrations like copepods and krill (Forward, 1988; Mauchline, 1998), suggests common biophysical mechanisms that drive evolution, fitness, and ecology.

Coordinating several appendages metachronally rather than synchronously induces steadier flow to generate thrust more efficiently (Ford et al., 2019). The antiplectic metachronal wave – propagating in the swimming direction – causes anterior appendages to actuate after their posterior counterparts and promote beneficial tip vortex interactions (Ford and Santhanakrishnan, 2021). The induced wake is opposite to the swimming direction and can interact constructively with the downstream appendages to further enhance thrust (Ford and Santhanakrishnan, 2021; Garayev and Murphy, 2021). These mechanisms are particularly sensitive to appendage synchronicity (i.e., phase) and spacing (Alben et al., 2010; Ford and Santhanakrishnan, 2021; Ford et al., 2019; Garayev and Murphy, 2021) for which most metachronal swimmers fall within a narrow range (reviewed in (Byron et al., 2021) Metachronal appendages must also achieve significant spatial or temporal asymmetry between the power and recovery strokes to maximize thrust production (Herrera-Amaya and Byron, 2023; Lou et al., 2022).

The biophysical commonalities among metachronal animals have been consistently found to promote the highest average flux and fluid momentum, thereby maximizing performance during forward swimming (Alben et al., 2010; Byron et al., 2021; Ford and Santhanakrishnan, 2021; Ford et al., 2019; Herrera-Amaya and Byron, 2023; Khaderi et al., 2012). As such, metachronal propulsion has been thought of as a strategy whose performance stems from maximizing thrust production (Alben et al., 2010; Byron et al., 2021; Colin et al., 2020; Ford et al., 2019). However, other major commonalities at the appendage and whole-system levels can be equally crucial in reducing drag, which is another critical performance factor to consider in self-propelled systems.

At large scales, crustaceans have complex biramous swimming legs (pleopods) and fine setae that open and close during the power and recovery phases, respectively. Recent studies have highlighted the effect of dynamically modulating the effective surface area of the propulsors (the projected area perpendicular to the flow) in producing spatial asymmetry between the power and recovery strokes (Byron et al., 2021; Herrera-Amaya and Byron, 2023; Lou et al., 2022; Vogel, 2020). While this has been shown to effectively overcome both the resisting body drag and the drag on the appendages to achieve propulsion, the role of bending has yet to be evaluated.

Metachronal swimmers keep their appendages relatively stiff during the power stroke but display significant spanwise bending to reduce the effective profile area, and thus any form drag, during the recovery stroke (Hessler, 1985; Johnson and Tarling, 2008). Ciliated microorganisms generate asymmetric waveforms by actively sliding adjacent microtubules within the propulsor (Dutcher, 2019). However, whether other larger metachronal organisms like crustaceans achieve active appendage spatial asymmetry using a similar mechanism is unknown. Passive asymmetrical bending resulting from the inherent structural or material properties of the appendages is more likely given the high mechanical stability and functional adaptation of crustacean exoskeletal elements through structural and compositional diversity (Fabritius et al., 2016; Raabe et al., 2005). By bending under fluid loading, flexible appendages can reduce their effective surface area and become more streamlined (Vogel, 2020). Self-reconfiguration not only lowers the drag coefficient but also reduces the stress on the structure proportionally to the resistive forces of the fluid (Alben et al., 2002; Gosselin et al., 2010).

Most metachronal swimmers produce antiplectic waves (Byron et al., 2021; Colin et al., 2010; Daniels et al., 2021; Garayev and Murphy, 2021; Murphy et al., 2011; Sensenig et al., 2010; Tamm, 2014) that cause two or more adjacent appendages to coalesce such that their profiles overlap. This is made possible by the proximity and relative phases of neighboring appendages (Byron et al., 2021; Murphy et al., 2011). As posterior appendages enter their recovery phase, they press against the anterior appendages that are transitioning from the power to the recovery stroke. This is particularly apparent in metachronal animals operating in the transitional to inertial flow regimes with Reynolds number, Re > 10 (Blake and Sleigh, 1974; Daniels et al., 2021; Garayev and Murphy, 2021; Murphy et al., 2011).

Coalescence may leverage the interactions between multiple flexible appendages to further reduce the total effective surface area to the flow, thereby lessening interactions with the fluid and lowering drag during the appendage recovery stroke. While stiff appendages can still coalesce, by virtue of proximity and phase, interactions within a group might cause destructive collisions or incomplete contact causing flow instabilities. Flexibility presents distinct advantages in this context.

Appendage asymmetrical bending and coalescence are apparent from simple observations of freely swimming animals. However, we still need a systematic understanding of the relationship between flexibility, temporal coordination, and coalescence to explain how these mechanisms reduce drag to enhance propulsion. Recent computer fluid dynamics (CFD) models provided insights into how metachronally beating appendages exploit a vortex-weakening mechanism to produce less overall drag and improve the thrust-to-power ratio significantly (Lionetti et al., 2023). However, experimental data about the underlying biophysical principles of drag reduction in metachronal propulsion are scarce. It is experimentally challenging to measure the flow among closely spaced appendages and disentangle the resulting forces. Still, several biological and robotic studies successfully identified how krill generate thrust and lift by estimating the momentum of the wake (Ford and Santhanakrishnan, 2021; Ford et al., 2019; Murphy et al., 2013). Yet, the contributions from individual and interacting legs and the independent effects of appendage bending and coalescence on the swimming performance could not be isolated. In addition, the recurrent simplification of appendage morphology as rigid rather than flexible plates (Ford and Santhanakrishnan, 2021; Murphy et al., 2011; Ruszczyk et al., 2022) might have also inadvertently undermined the significance of appendage bending as a major contributor to drag reduction.

This investigation presents a comprehensive overview of the morphological, functional, and physical mechanisms promoting drag reduction during metachronal swimming by establishing the relationship between appendage flexibility, temporal coordination, and coalescence. Using μ-CT measurements, we examined the morphology of marsh grass shrimp (*P. vulgaris*) pleopods to explore the characteristics enabling passive asymmetrical bending. We combined free-swimming, *in-vivo* kinematics and velocimetry experiments to quantify bending and coalescence and visualize their hydrodynamics. Using this biological data and leveraging our Pleobot (Santos et al., 2023), we designed a scaled metachronal robot with five morphologically accurate and flexible pleopods capable of asymmetrical bending. We quantified the thrust and drag of independent and coalescing pleopods and evaluated the resulting wake using simultaneous force and fluid flow measurements. We compared these results against rigid pleopods to answer the following three questions: Does pleopod bending significantly reduce drag during the leg recovery phase? Given that shrimp group three recovering pleopods, does coalescence turn the drag of several appendages to that of only one? How complementary are bending and coalescence in optimizing drag reduction? We provide evidence that reducing drag, rather than simply enhancing thrust, is a determinant factor in the functional morphology and coordination of metachronal propulsors. Our results leverage our understanding of the unifying biophysical principles of this locomotor mode that can be applied to the design of novel bioinspired underwater robots.

## MATERIALS AND METHODS

### Animals

Marsh grass shrimp (*P. vulgaris*) (n = 13; body length = 3.11 ± 0.25 cm) were collected in June 2022 from Narragansett Bay (Rocky Point State Park, Warwick, RI, USA). The shrimp were housed at room temperature (21°C) in a 38-liter aquarium. The capture and experimentations were conducted in accordance with the laws of the State of Rhode Island.

### Micro CT scan

A shrimp specimen was euthanized in 25% ethanol (seawater, 30 ppt) and transferred to a 70% ethanol solution for 20 min until rigor mortis occurred. The left and right pleopods (from P1 to P5) were amputated at the proximal end of the protopod. This was done to separate the pleopods to avoid physical interactions and to enhance the clarity of the resulting high-resolution μ-CT scan. The samples were preserved in 100% ethanol for five days and stained in a 1% solution of Iodine in 100% ethanol for 24 h. They were then submerged in 98% Hexamethyldisilazane (HMDS) for 24 h and air-dried overnight. Chemical drying using HMDS produces similar results to critical-point drying but allows larger samples to be processed (Alba-Tercedor and Alba-Alejandre, 2019; Bhattacharya et al., 2020). The samples were finally mounted onto a custom-made 3D-printed mount that fit the holding tray of the μ-CT scanner. A SkyScan 1276 high-resolution microtomography (Bruker, Billerica, MA, USA), upgraded to receive a Hamamatsu microfocus X-ray source (L10321-67, Bridgewater Township, NJ, USA) and a 4K camera (XIMEA MH110XC, Lakewood, CO, USA) was used to perform high-resolution scans (see supplemental materials for specific scanning parameters). We used the Bruker micro-CT’s Skyscan software (NRecon, DataViewer, CTAnalyser) for primary 3D reconstructions from cross-section images. Volume rendering images were obtained with Slicer v.5.0.3. The 3D reconstruction was used to extract transverse sections of the endopodite and exopodite of the ten legs of the specimen (pooling left and right appendages) to measure the radius of curvature of the chord, *R*, giving the chordwise curvature *κ* as its reciprocal (see details in the supplementary materials). Curvature was measured from chordwise cross sections halfway through the length of each ramus. To account for differences in the width of the ramal structures that proportionally impact the raw value of the radius of curvature, we normalized *κ* to the width, *W* of the ramal structures (established from the above transverse sections) as *κ = W/R*. Note that we excluded the chordwise curvature of the left and right P1 endopodites from analyses because they are sexually dimorphic, atrophied, and not involved in propulsion.

### Pleopod flexural stiffness

Five specimens were euthanized and all their pleopods were amputated at the proximal end of the protopod and were used within one hour. The fresh protopod of each pleopod was fixed to a rigid metal pin using cyanoacrylate glue by immobilizing the peduncle-ramal joint. In all cases, either the endopodite or the exopodite of one leg in a pair was randomly amputated to avoid mechanical interactions during testing. The second ramal structure was isolated similarly using the opposing leg. The mounted pleopods were immersed in 30‰ salt water to maintain the natural material properties and were placed on a precision balance (VWR-124B2, VWR International, USA, precision = 0.1mg). We applied a series of point force loads at the appendage tip on the ventral and dorsal sides, corresponding to the power and recovery strokes, respectively. Specifically, we applied the point force at the tip of the membranous part of the rami rather than the tip of the projecting distal setae. This is because the setae have different material properties and spread out, thus letting the pin through. Five replicates were obtained on both sides of each structure and two pictures were taken for each replicate, starting with the unloaded state and then the loaded state. We analyzed each image of the deflected pleopod unit using MATLAB R2022a (MathWorks, Natick, MA, USA) to determine its effective beam length and deflection. Beam length (*L*, in m) was measured as the distance from the peduncle-ramal joint to the point of applied force. Deflection (δ, in m) was measured as the mean vertical distance traveled by the tip of the appendage from its initial reference location for each set of five replicates. Overall spanwise flexural stiffness *EI* was measured using the beam equation as:

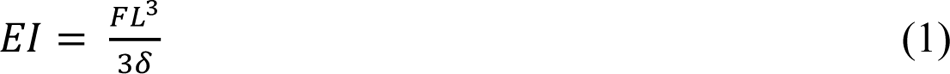

where *F* is the product of the mass reported on the balance readout and the gravitational acceleration (in Newtons). This equation measures the flexural stiffness over the entire beam length and assumes that the beam is homogeneous. Because the equation applies only to small displacements, we removed any measurements for pleopod displacement δ > 0.08*L*. This is because if δ is large relative to *L*, the beam is bent down so far that its coordinates in the x-dimension while loaded are significantly different from its coordinates while unloaded, thus invalidating measurements(Combes and Daniel, 2003). Eq. 1 is valid for our measurements because even in the most extreme cases (where δ is near 0.08*L*), the error in coordinate introduced by length changes in the x-dimension was less than 0.3% (equivalent to the measurement error).

### Flow visualization

Free-swimming experiments were performed in a 9.5-l aquarium. Individual shrimp were brought near the center of the tank upstream of the field of view and within the focal plane of the camera using a beaker. After the flow stabilized, the shrimp were allowed to exit the beaker and swim freely through the field of view, reaching constant speed by the time they entered it. We performed bright-field PIV by seeding the water with 10 μm particles (Dantec Dynamics, Skovlunde, Denmark) and back-lighting the animals with an LED illuminator (M530L2-C3, Thor Labs, Newton, NJ, USA) coupled with a collimating Fresnel lens (focal distance = 10 cm). We recorded images from the lateral view (corresponding to the left side of the animals) using a high-speed digital video camera (Fastcam Nova R2, Photron, Tokyo, Japan) at 2000 frames s^−1^ at a resolution of 2048 × 1472 pixels. The camera was fitted with a 60 mm macro lens (Nikon, AF Micro Nikkor, Nikon, Japan) set at f/5.6, yielding a depth of focus for PIV analyses DoF ≈ 57.1 μm (see supplementary materials). Fluid velocity vectors were calculated from sequential images analyzed using the DaVis 10 software package (LaVision, Göttingen, Germany). Image pairs were analyzed with three passes of overlapping interrogation windows (75%) of decreasing size of 64 × 64 pixels to 48 × 48 pixels. All frames were used for the analysis, yielding a separation between frames (dt) of 5 × 10^−4^ s. Velocity fields were input to a custom program in MATLAB 2022a (MathWorks, Natick, MA, USA) that computed the corresponding vorticity fields.

### Kinematics and morphometrics

Pleopod kinematics data were extracted from image sequences using a custom program in MATLAB. The profiles of individual pleopods (from P1 to P5) of the left side of each shrimp were digitized by manually fitting a Bezier curve controlled by two handles. This method accurately matches the deformation of the pleopods during both the power and recovery stroke. Because the protopod is rigid, it was digitized as a straight segment, while the endopodite was digitized as a Bezier curve. We only digitized the endopodite because the exopodite abducts during the power stroke and rotates out of plane. In contrast, the endopodite remains aligned with the focal plane. Every 10 frames were used for analysis, yielding a separation between frames (dt) of 5 × 10^−3^ s. Pleopod spanwise bending during swimming was quantified using the curvature along the ramal structures for each leg. Points along the pleopod of length *L* were specified as fractional distance from the peduncle-ramal joint. The radius of curvature *R* was measured at all points along the pleopod Bezier profiles. We used curvature, *κ* (the reciprocal of *R*), normalized to pleopod length *L*.

The α angle, defining the pleopod beat amplitude, was calculated by measuring the angle between the protopod and the line passing through the proximal joint of the protopods of P1 and P5 (aligned in the direction of swimming). The β angle is traditionally measured between the protopodite and the line passing through the proximal and distal sections of the ramal structures. However, because the endopodite and exopodites bend, we instead measured β between the protopod and the line tangent to the proximal section of the endopodite (corresponding to 20% of the total length of the endopodite) to account only for the movement due to the peduncle-ramal joint. A Fast Fourier transform of the α angle over time was used to determine the sequence-averaged pleopod beat frequency.

Pleopod coalescence was quantified through the calescence ratio (CR) for each pleopod as the ratio of pleopod grouping duration during a beat (when in contact with at least one adjacent pleopod) to the duration of the corresponding recovery stroke. Coalescence was discretized using three categories: leading (when the pleopod in a coalescing group is facing the flow), confined (when the pleopod is surrounded by an anterior and posterior pleopod), and trailing (when the pleopod in a group is posterior the others). We defined the metachronal wave period (T_wave_) as the time between the beginning of the power stroke of P5 (initiating the wave) and the end of the recovery stroke of P1 (ending the wave), after all the other pleopods have completed their cycle. For P2–P4 with anterior and posterior neighbors, coalescence starts when a recovering posterior pleopod contacts the posterior face, followed by an anterior pleopod contacts the anterior face upon finishing its power stroke. The phase φ between two consecutive pleopods was measured as the time between the start of the power stroke of a corresponding pleopod relative to that of its posterior neighbor (the latter propagating the metachronal wave earlier) and was normalized to its beat period (calculated as 1/*f* where *f* is the beat frequency). The ratio of pleopod spacing (*B*) to pleopod length (*L*) was calculated as the distance between the proximal protopodite joint of a corresponding pleopod and that of its posterior neighbor, divided by the sum of its protopodite and exopodite lengths. Because P5 does not have a posterior neighbor, *B*_P5_ was measured as the distance between P4 and P5.

### Robotic analog

We used the mean leg morphometrics (n = 10 shrimp) and kinematics data for the α and β angles of one representative shrimp to design and operate a 20× scaled 3D printed five-legged shrimp robotic analog (Fig. 1, Table 1). The kinematics data from a single specimen demonstrating horizontal steady swimming was used to drive the robotic leg motions with no destructive interference between pleopods, as averaging data from multiple specimens masked individual variations and caused physical contact issues (see supplementary materials). The design and actuation of the legs are based on the *Pleobot* (Santos et al., 2023), whose kinematics and hydrodynamics have been validated experimentally against biological data. Here, we fitted the robot with flexible ramal structures (Practi-Shim™, thickness = 0.0254 mm), whose chord profile was heat-shaped using the same curvature profile of shrimp pleopods to achieve differential stiffness (see supplementary methods). Note that because the P1 endopodite is sexually dimorphic, atrophied, and not involved in propulsion in *P. vulgaris*, the robotic P1 pleopod did not include an endopodite. The viscosity of the fluid and the beat frequency of the legs were adjusted to match the Re of the appendages of the five-legged robot to that of the marsh grass shrimp used in in-vivo experiments (Re_robot_ ≈ Re_shrimp_ = 1720, see supplementary materials). A 3:2 glycerin-water mix with density of ρ = 1160 kg m^−3^ and dynamic viscosity, μ = 0.0114 kg m^−1^ s^−1^, was used in the experiments.

**Figure 1.**
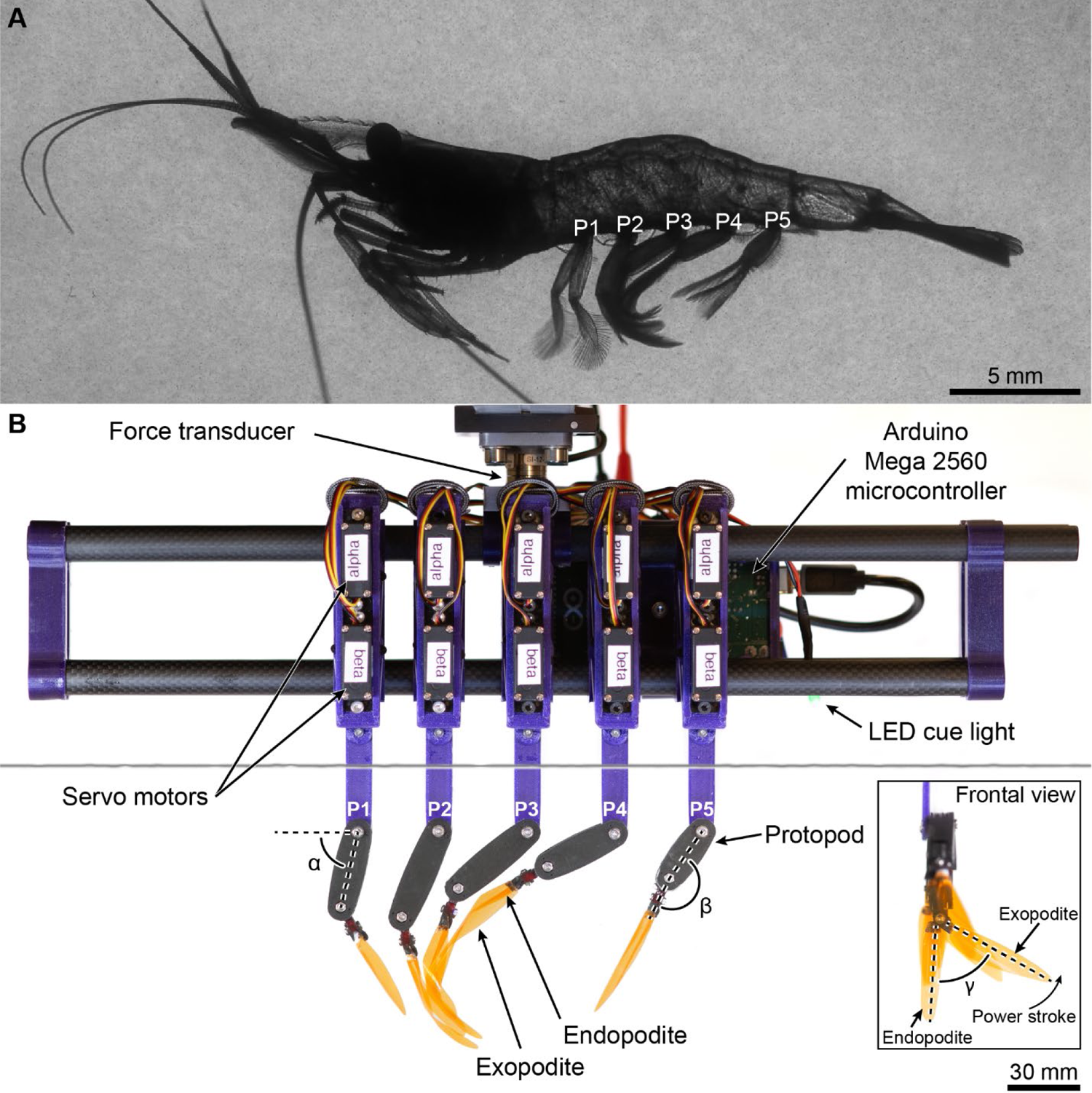
Biological and robotic models. **(A)** Representative free-swimming marsh grass shrimp (*P. vulgaris*). **(B)** The tethered bioinspired 20× robot with five flexible appendages replicates pleopod bending and coalescence, and measures forces. We used pleopods representative of the left side of the animal, consistent with kinematics data of free-swimming shrimp. Pleopods are identified from P1 (anterior) to P5 (posterior). The α, β, and γ angles are shown for reference. The horizontal gray line is the water surface.

**Table 1.**
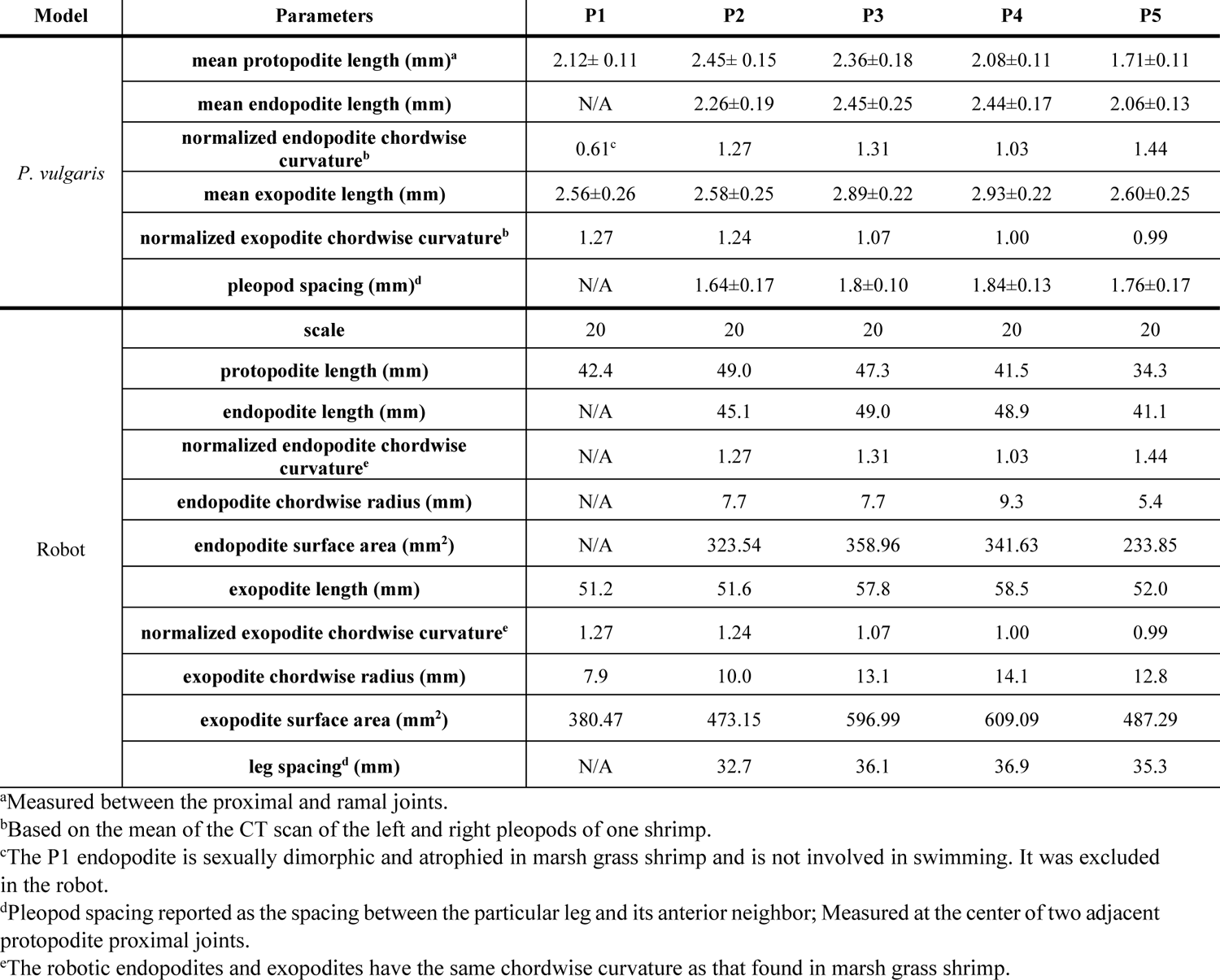
Comparative leg morphometrics of *P. vulgaris* and the robotic analog. Mean protopodite, endopodite, and exopodite lengths for *P. vulgaris* are reported as mean ± s.d. (n = 10 shrimp).

We performed high-speed 2D PIV and force measurements synchronously to temporally match the kinematics and PIV sequences to the resulting instantaneous forces produced by and acting upon the pleopods. Measurements were acquired after at least ten consecutive metachronal waves to ensure the flow was fully established. Forces were measured by a 6-axis Nano 17 force transducer (ATI Industrial Automation, MI, USA) from which the robot was suspended. The transducer was interfaced to a data acquisition (DAQ) system (NI USB-6210, National Instrument, USA, TX) outputting data to a custom MATLAB script that converted raw voltages to force data. The sensor bias was measured before each experimental trial and subtracted from the force data to eliminate the potential effects of sensor drift over time. The robot was initially tested in air to measure the inertial effects during leg motions to account for the added mass during the experiments. However, because of the low beat frequency and the fact that the pleopods have low mass by virtue of being made of thin plastic shims, the forces due to inertia were too small to be distinguishable from the static noise produced by the force transducer under its minimum measurement threshold (uncertainty = 2.25 mN). Therefore, we considered added mass effects to be negligible in the experiments.

We used high-speed two-dimensional PIV to compute velocity fields around the robotic legs. Experiments were carried out in a 210-liter aquarium measuring 119 × 43 × 41 cm (internal length × width × height). The robot was suspended above the surface of the fluid with its pleopods fully submerged. Lateral recordings were acquired by a high-speed, high-resolution digital video camera (Fastcam Nova R3, Photron, Tokyo, Japan) at 500 or 750 frames per second (4096 pixels × 2304 pixels). Seeding particles (10 μm Dantec Dynamics, Skovlunde, Denmark) were illuminated by two adjacent coplanar vertical laser sheets (532 nm, 4000 mW, OptoEngine LLC, Optotronics LLC, Mead, CO, USA) to ensure a uniform particle density field along the entire length of the robot. Both laser sheets pointed upward along the medial plane of the robot (vertical plane passing through the tips of the endopodites).

Several leg configurations were tested to isolate the effects pleopod asymmetrical bending and coalescence have on the fluid flows and forces. For instance, we tested individual pleopods (P1– P5) fitted with flexible or stiff appendages to determine the thrust and drag generated in both cases. Each pleopod was tested independently by keeping all the other legs horizontally, out of the way and influence of the flow produced by the selected pleopod. To investigate the effects of coalescence, we performed five-legged and three-legged experiments. The five-legged cases were analogous to the free-swimming marsh grass shrimp while the three-legged cases aimed at precisely reproducing a unique coalescing group of three adjacent pleopods. This was done to eliminate the effects of the other legs not involved in the corresponding coalescing group. We report data for the P2-P3-P4 coalescing group. To test the effect of coalescence on the flow and forces specifically, we compared the results to the instantaneous forces produced by the same, but independent, legs, time-synchronized to the corresponding metachronal wave of the coalescing group. The theoretical thrust when coalescence is not occurring was calculated as the cumulative axial force produced by the individual legs. We also tested cases when the metachronal wave was phase-shifted by 180 degrees. This unnatural kinematics caused leg contacts during the power stroke, but complete separation of the pleopods during the recovery stroke. Lastly, to explore the compounded effect of bending and coalescence, we tested the above leg configurations with flexible and stiff ramal appendages. Fluid velocity fields were computed using the DaVis 10.2 software package (LaVision, Göttingen, Germany). Image pairs were analyzed with three passes of overlapping interrogation windows (75%) with decreasing size from 64 × 64 pixels to 48 × 48 pixels. No smoothing of the velocity data was performed.

The force coefficient *Cf*, whose thrust and drag coefficients (*Ct* and *Cd*, respectively) were of interest in this investigation, was calculated using the axial forces oriented in the swimming direction, corresponding here with the body axis of the five-legged robot. In this context, the axial forces generated by individual legs contribute entirely to thrust (in the swimming direction) or drag (opposite to the swimming direction). The cycle-averaged coefficients were computed over 8–10 consecutive cycles (based on availability of complete cycles at the beginning and end of the video sequence) from the instantaneous force magnitude *T*, as:

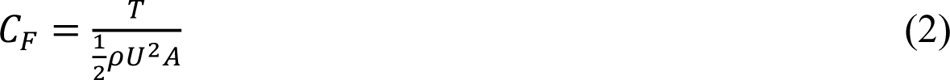

where *ρ* is the density of the 3:2 glycerin-water mix, *U* is the cycle-average appendage tip velocity, and *A* is the stroke-phase dependent total surface area of the appendage. *A* was maximal during the power stroke as the exopodite extended fully (see Fig. 1), thus combining the surface area of the endopodite and exopodite in the calculation of the force coefficients during the power stroke. During the recovery stroke when the exopodite adducts and overlaps completely with the smaller endopodite, the total surface area of the pleopod is reduced to the surface of the exopodite only. The coefficients are calculated using the total surface area of the pleopods rather than their effective surface area – as if both pleopod types were stiff. This is to highlight the expected difference in the drag the stiff and bending pleopods produce (due to different effective surface area) despite having the same kinematics, shape, size and total surface area. *U* was calculated assuming the tip of the appendage moves along an arc along the imaging plane during a beat as:

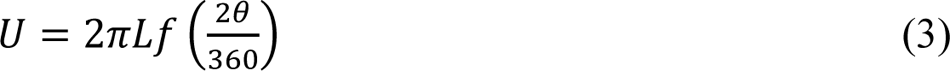

where *L* is the total length of the pleopod (including protopodite and exopodite), *f* is the beat frequency, and *θ* is the appendage stroke amplitude, in degrees, obtained from the α angle maxima and minima over several consecutive beats. The α angle was measured for each leg relative to the swimming direction by measuring the leg kinematics using the DLTdv8 package for MATLAB. This was done by tracking landmarks on the protopodite, such as the distal joint screw. The slight orientation bias of the rostro-caudal axis relative to horizontal (bias < 1°) was accounted for in the α computation.

### Statistics

All statistical tests were performed using MATLAB 2023b. For biological measurements of the pleopod chordwise curvature, the rami were pooled by type for the only specimen available (for a total of 8 endopodites and 10 exopodite). Note that the sexually dimorphic and atrophied P1 endopodite were excluded from this analysis because they differ substantially from the other legs and are not involved in swimming. Because of the low sample count and the sample distributions were not normally distributed (One-sample Kolmogorov-Smirnov test, *p*-value < 0.001), we compared the differences between the endopodite and exopodite using a Mann-Whitney *U*-test. Similarly, the flexural stiffness of the anterior and posterior faces of the endopodite and exopodite (n = 5 shrimp), was compared using a *U*-test for each leg. The relationship between pleopod bending and the angular velocity ω of the pleopod defining its overall motion relative to the fluid was examined separately for the power and recovery strokes. For the recovery stroke, we found that the relationship between the local curvature *κ*/*L* along the ramus of length *L* pleopod angular velocity is best described by a power fit (*κ*/*L* = aω^b^ + c) while the power stroke is best modeled as a linear fit (*κ*/*L* = aω^b^ + c). For robotics experiments, the mean thrust and drag coefficients for each consecutive beat cycle (for individual legs fitted with flexible or stiff ramal appendages) were computed for the power and recovery strokes, respectively, to calculate the cycle-average mean coefficients (from 8 to 10 consecutive cycles). We performed *t*-tests to compare the mean thrust and drag coefficients of individual flexible pleopods with their stiff counterparts (P1 through P5). Differences were considered significant at *p*-values < 0.05. All the kinematics parameters and force data are reported as mean ± standard deviation (s.d.).

## RESULTS

### Pleopod morphology and asymmetrical bending

Cross sections of all the pleopods of one representative *P. vulgaris* obtained from reconstructed μ-CT scans show that the posterior face of the ramal structures — endopodite and exopodite — is concave, with pronounced chordwise curvature (Fig. 2A). Pooling the data by ramus type revealed no statistical difference in the normalized curvature *W/R* of the endopodites and exopodites (8 endopodites and 10 exopodites, *U*-test, *p* = 0.083, Fig. 2) with *W/R*_endopodite_ = 1.26 ± 0.21 and *W/R*_exopodite_ = 1.11 ± 0.14 (see Table 1 for *W/R* of individual pleopod). Note that the chordwise curvature of the left and right P1 endopodites were excluded from analysis because they are sexually dimorphic, atrophied, and not involved in propulsion. We observed that the anterior face of the endopodites and exopodites was relatively planar and determined that chordwise curvature was the feature enabling asymmetrical bending. Hence, we simplified the model by only considering the curved profile in the design of the robotic appendages. The rami of each robotic pleopod were shaped using the normalized curvature of the rami of the respective leg measured in *P. vulgaris* (Table 1).

**Figure 2.**
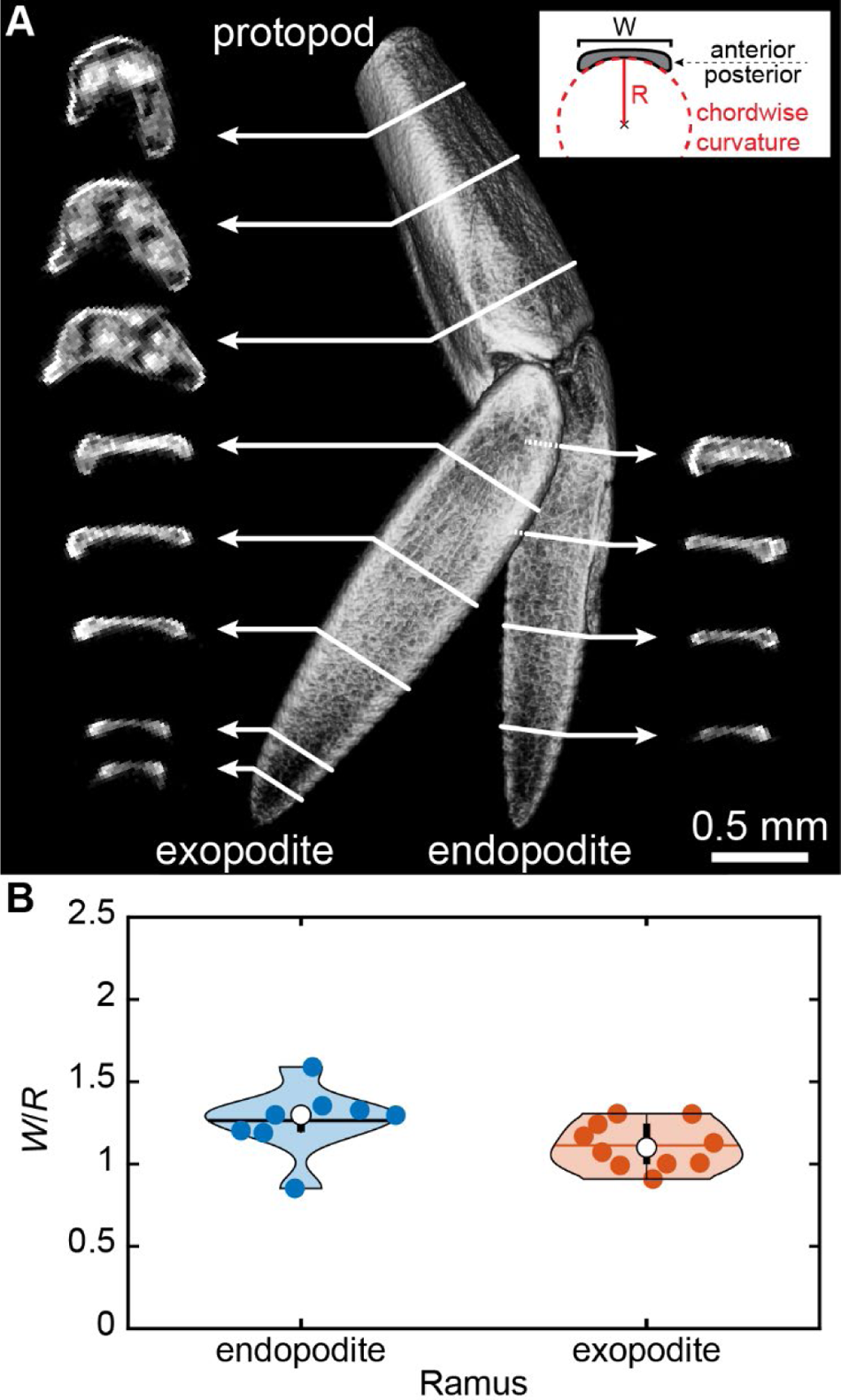
Chordwise curvature of the pleopods. **(A)** Reconstructed μ-CT scan of the right P2 pleopod of a representative *P. vulgaris* specimen (anterior view). Cross sections show the chordwise curvature, *W/R* of the posterior face of the exopodite and endopodite, with *W* as the width and R the radius of curvature (see inset). Note the flat anterior face. **(B)** Corresponding mean normalized chordwise curvature of the posterior face of the endopodites and exopodites (n = 10 legs). Endopodites and exopodites have comparable chordwise curvature (*U*-test, *p* = 0.083).

The posterior chordwise curvature of the rami enabled the legs to remain stiff during the power stroke and bend almost horizontally during the recovery stroke (Fig. 3A). Flexural stiffness measurements of freshly dissected *P. vulgaris* pleopods show the rami are, on average, 1.7 times stiffer when a point force is applied posteriorly (corresponding to the power stroke) compared with the anterior face (analogous to the recovery stroke), resulting in pronounced spatial asymmetry (*U*-tests, 0.008 < *p* < 0.343; n = 5 shrimp; Fig. 3, Table S1). Stiffness measurements were obtained for the membranous part of the rami and excluded the setae.

**Figure 3.**
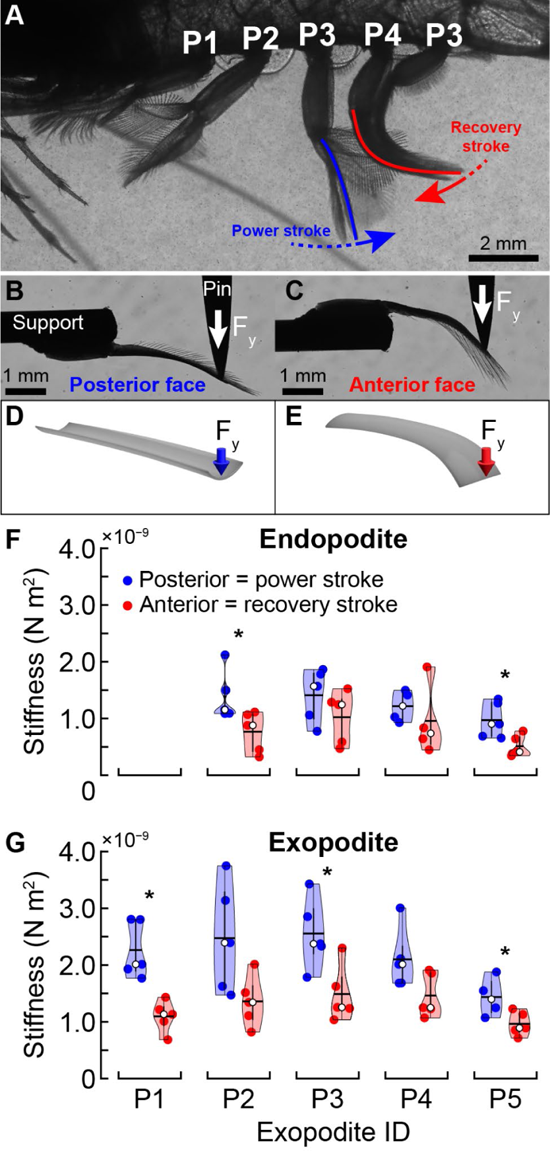
Differential flexural stiffness of *P. vulgaris* pleopods. **(A)** Differential stiffness keeps the pleopods stiff during the power stroke and allows bending during the recovery stroke. **(B)** Posterior and **(C)** anterior measurements of pleopod flexural stiffness. Pleopod deflection is emphasized to illustrate differential stiffness. **(D)** Conceptually, the concave posterior face stiffens the rami when a force is applied posteriorly (power stroke) while an anterior force causes bending **(E)**. The endopodites **(F)** and exopodites **(G)** were, on average, 1.7 times more flexible during the recovery vs. the power strokes (*U*-tests, 0.008 < *p* < 0.343; n = 5 shrimp). Asterisks indicate statistical differences *p* < 0.05 between the anterior and posterior faces. The white indicates the mean and the horizontal black line is the median.

Pronounced pleopod spanwise bending occurs exclusively during the recovery stroke, as seen with elevated local curvature propagating from the proximal to the distal sections of the rami (Fig. 4A, Fig. S4). The rami do not bend along a singular point. Instead, the inflection point induced in the proximal section at the beginning of the recovery stroke travels distally over time until the end of the recovery phase. The rami remain nearly horizontal during the first half of the recovery phase (during the acceleration phase of the recovering pleopod) but eventually unfurl completely in preparation for the following power stroke (Fig S5). We found that the rami bent passively — rather than actively — due to the resistive forces of the flow acting on the anterior face during the recovery stroke. We can infer the resistive effects of the flow on the pleopod via the instantaneous angular velocity, as in this regime (Re ≈ 1700) the drag scales with the square of the velocity (the linear velocity at any point along the span of the rami) of the appendage. By measuring the instantaneous local curvature at a fixed point along the rami during a beat (we selected 0.3*L*, n = 6 shrimp), we found that spanwise bending of the rami was proportional to the angular velocity of the pleopod. It was particularly significant during the recovery stroke when the local curvature scaled as a power relationship with pleopod angular velocity (*κ*/*L* = aω^b^ + c, Fig. 4B, Fig. S4). In contrast, the rami bent only very little with increasing pleopod angular velocity during the power stroke, which followed a linear trend in this case (*κ*/*L* = aω + b, Fig. 4B, Fig. S4). This is because the pleopod must remain stiff to generate maximum thrust.

**Figure 4.**
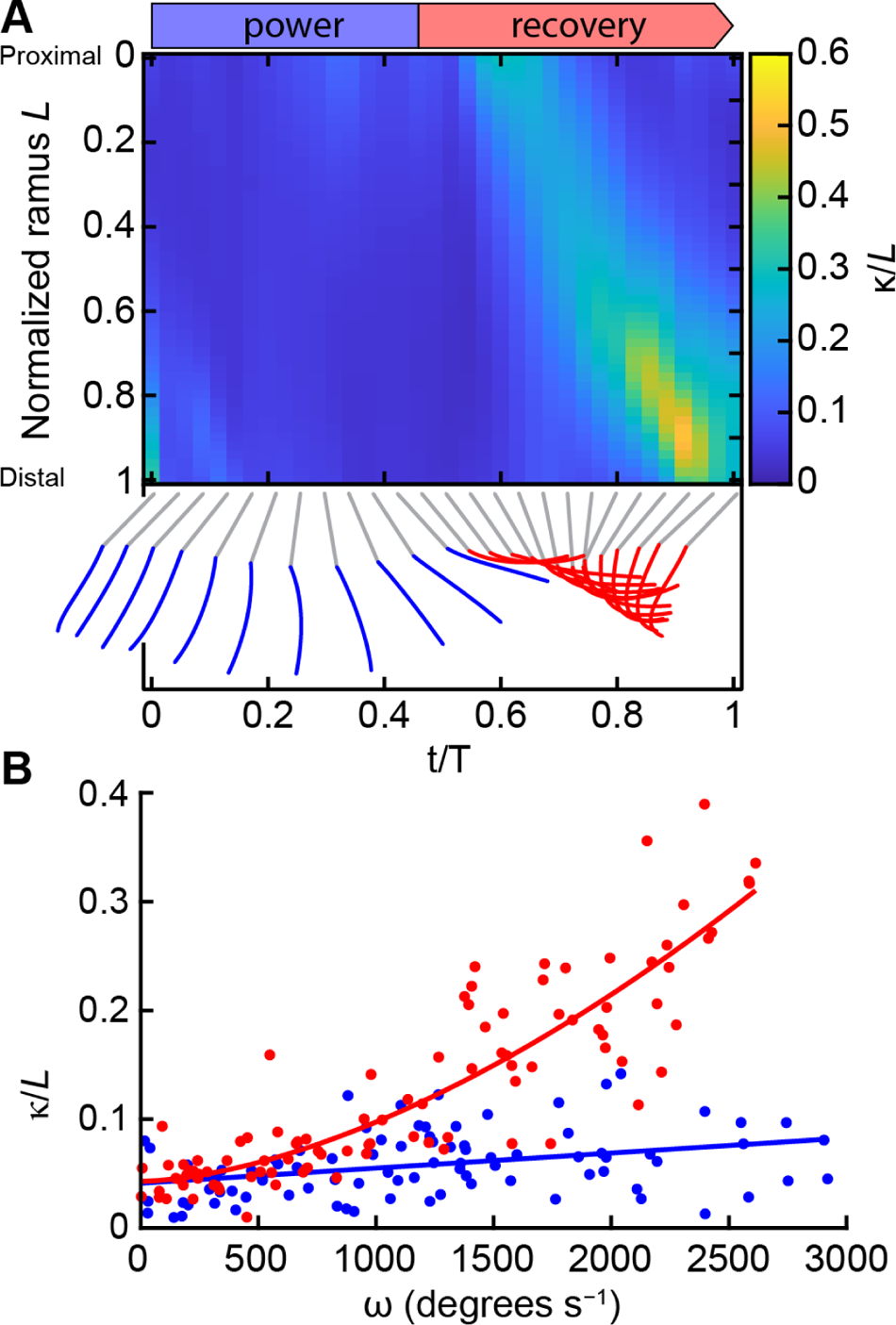
Passive pleopod asymmetrical bending during a beat. **(A)** Local normalized curvature along the flexible P3 exopodite during a full beat cycle. Instantaneous pleopod profiles (blue and red) highlight differences in curvature between the power and recovery strokes. **(B)** Local spanwise curvature at 0.3*L* of P3 as a function of angular velocity during a full beat (n = 6 shrimp). The local curvature increased with the angular velocity (ω) of the pleopod during the recovery stroke following a power relationship (*κ/L* = 5.84×10^−7^ω^1.66^ + 0.04, R^2^ = 0.78). The power stroke induced little bending and was best modeled using a linear fit (*κ/L* = 1.39×10^−5^ω + 0.04, R^2^ = 0.122).

Individual, isolated robotic pleopods fitted with flexible rami behaved similarly and had comparable fluid-structure interactions (FSI) to their rigid counterparts during the power stroke (Fig. 5A,C). In both cases, the pleopods remained stiff to sustain the largest surface area and generate maximum thrust. For each pleopod, flexible and stiff rami generated identical power stroke (PS) tip vortices and had equivalent mean thrust coefficients (*t*-tests, 0.09 < *p* < 0.58, Fig. 5E,F). Both cases shed the PS tip vortex from the pleopod tip (Fig. 5B,D). The most notable differences between the flexible and stiff cases emerged during the recovery stroke as the flexible rami allowed the PS wake to circulate posteriorly unimpeded (Fig. 5B). In contrast, the stiff rami prevented most of the PS wake from circulating rearward, causing a significant increase in the drag experienced by the leg. By virtue of bending nearly horizontally, the flexible appendages interacted only marginally with the PS wake and had a reduced effective surface area to the flow compared to the stiff rami. This resulted in a significant reduction of the drag coefficient *C_D_* of P2 through P5 by 41.0, 66.4, 75.8, and 42.6% respectively (*t*-tests, *p* < 0.05 for P2–P5 and *p* = 0.08 for P1; Fig. 5F). While the flexible rami induced a dominantly rearward wake, the stiff rami shed a recovery stroke (RS) tip vortex carrying forward momentum (Fig. 5C). We did not find evidence that these flow-induced FSI impacted either the anterior flow along the pleopod or the thrust coefficient significantly. Note that all the pleopods have a positive *C_F_* during the second half of the RS, which corresponds to the deceleration phase of the stroke. Additionally, unlike the posterior pleopods, P1 and P2 finish and initiate a beat with a significant positive *C_F_*. This is largely because of the effects of the posterior thrust-producing stagnating flow and the drag-inducing role of the interactions with the PS wake and induced flow. Because P1 and P2 have a smaller maximum α (α < 90°) than P3–P5 they produce a wake with a less pronounced posterior component which is largely overcome by the forces induced by the posterior stagnating flow. Note how the initial CF converges to 0 from the anterior to the posterior pleopods as their respective wake and induced flow becomes progressively more horizontal. While the flexible rami induced a dominantly rearward wake, the stiff rami shed a recovery stroke (RS) tip vortex carrying forward momentum (Fig. 5C). We did not find evidence that these flow-induced FSI impacted either the anterior flow along the pleopod or the thrust coefficient significantly.

**Figure 5:**
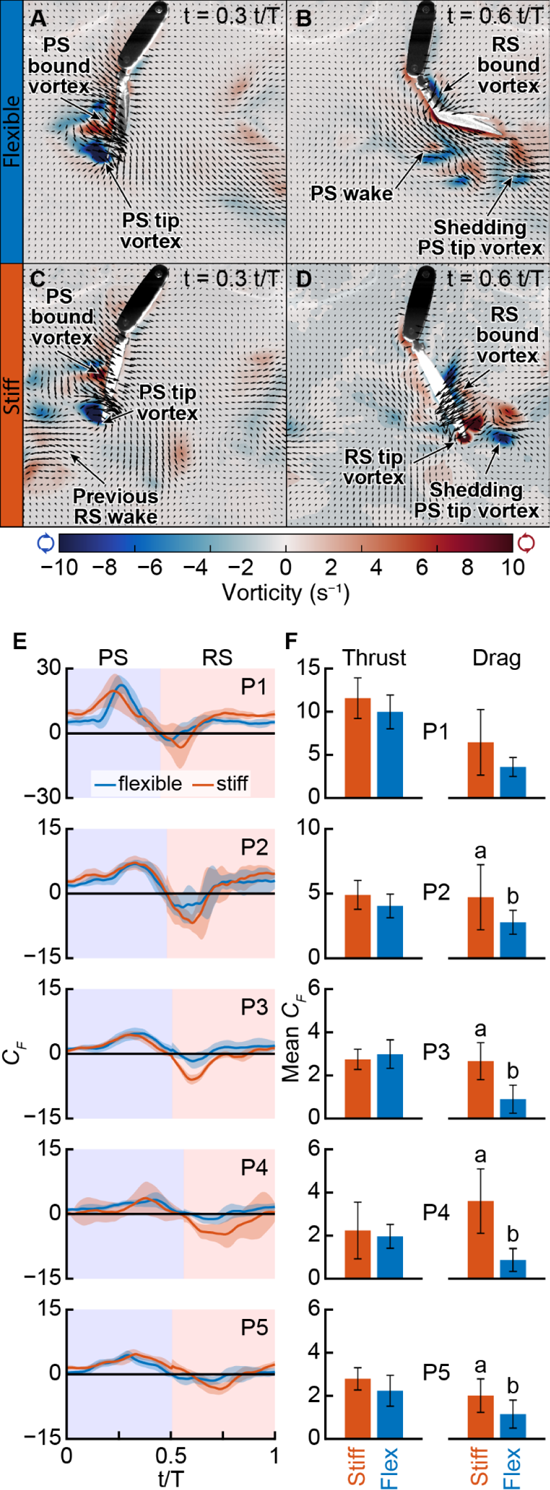
Flexible ramal structure generates less drag than their stiff counterparts. Instantaneous velocity and vorticity fields of P3 fitted with flexible ramal appendages **(A,B)** and stiff appendages **(C,D)**. Both appendage types generated comparable flow during the power stroke (PS) (**A,C)**. At the end of the recovery stroke (RS), the flexible rami induced a downward, posterior wake **(B)** while the stiff rami shed backward and forward wakes **(C,D)**. Every three vectors were plotted for clarity. **(E)** Cycle average force coefficient *C_F_* for flexible and stiff P1–P5 averaged for several consecutive beat cycles (n_flex_ ≤ 9 and n_stiff_ ≤ 10). The colored background indicates the PS (blue) and RS (red). Shading indicates the standard deviation. **(F)** Comparison of the mean *C_T_* (Thrust) and *C_D_* (Drag) of the flexible and stiff ramal appendages. Letters indicate significant differences between force types within a species (*t*-tests, *p* < 0.05).

### Pleopod coalescence

While pleopod bending plays a crucial role in decreasing the overall drag of each pleopod during the recovery stroke, it also enhances leg coalescence. Time series of coalescing pleopods in free-swimming *P. vulgaris* show that flexible pleopods can wrap around one another and conform to the shape of the surrounding pleopods to enhance coalescence (Fig. 6A, Fig. S5). Another benefit of flexibility is the significant reduction in potentially destructive physical interactions upon colliding with one another. Because of the length and proximity of the legs and the phase relative to one another, up to three pleopods form a tight coalescing group. Three-legged coalescence emerges from the specific pleopod-separation-to-length ratio, *B/L* = 0.364 ± 0.014 (n = 6 shrimp, all pleopods pooled) and inter-pleopod phase lag φ = 0.18 ± 0.02 t/T which causes several legs to perform their recovery strokes simultaneously (Fig. 6D).

**Figure 6.**
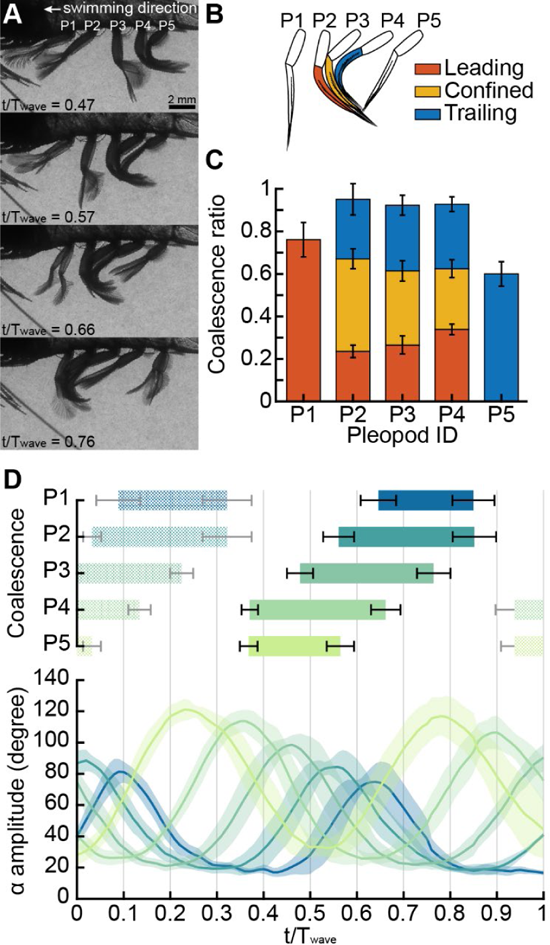
Pleopod coalescence. **(A)** Up to three adjacent pleopods coalesce during their recovery stroke (RS). **(B)** In a coalescing group, the anterior leg leads the movement while the posterior pleopod trails. The pleopod within a coalescing group is confined between the surrounding anterior and posterior legs. **(C)** The coalescence ratio defines the proportion of the RS during which pleopods coalesce. P2–P4 switched from leading to trailing the group (n = 6 shrimp). **(D)** Pleopod coalescence and kinematics visualized through the α amplitude during a full metachronal wave (n = 6 shrimp). The color code of the pleopod coalescence bars was applied to the α amplitude curves (from light to dark shades from P5 to P1 respectively). Errors bars and shading indicate the standard deviation for coalescence and α kinematics, respectively. Shaded coalescence bars indicate coalescence occurring during the previous and following metachronal waves.

The role of each leg changes during a coalescence cycle. For example, as initiates its coalescence cycle, it leads the P3-P4-P5 group and faces the flow directly. Upon reaching the anterior P2, it becomes confined between P2 (anterior) and P4 (posterior) which shields P3 from anterior and posterior flows. The coalescence group becomes P2-P3-P4. Finally, as P4 separates from the P2-P3-P4 group, P3 trails the P1-P2-P3 group, thus exposing it to posterior flow. P1 and P2 are exceptions since they only have one posterior and one anterior neighbor, respectively (Fig. 7).

**Figure 7.**
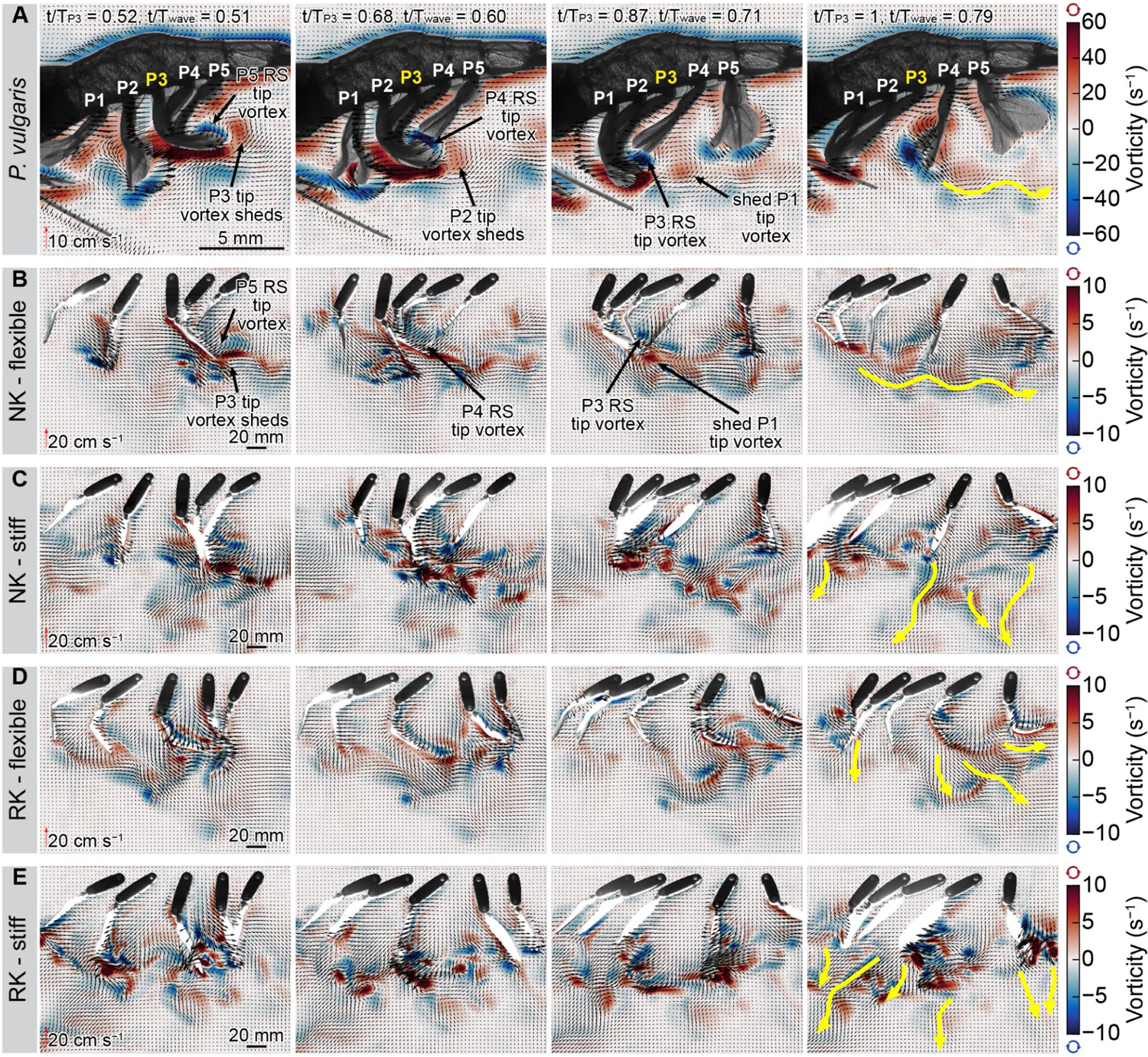
Velocity and vorticity field time series of *P. vulgaris* and the five-legged robot. **(A)** Free-swimming *P. vulgaris* displayed pleopod coalescence during the sequential recovery strokes (RS) of each leg. Note each pleopod shifting from leading to trailing the coalescing group as the metachronal wave propagates. **(B)** The five-legged robot with flexible rami and displaying natural kinematics (NK) generated a wake analogous to *P. vulgaris*. **(C)** Stiff rami produced a turbulent wake when the recovering pleopods intercepted the PS wake and generated individual wakes with forward and downward components. **(D)** Flexible rami performing reversed kinematics (RK) without coalescence also had a strong downward component. **(E)** Flexible, RK rami produced wakes with the most forward-oriented component out of all the above cases. Yellow arrows represent the overall direction of the pleopod wakes. For clarity, every two and three vectors are plotted for *P. vulgaris* and the robot, respectively.

We use the coalescence ratio (CR), to quantify the duration of leg coalescence for a given pleopod relative to the duration of its recovery stroke. P2, P3, and P4, which have one adjacent anterior and posterior pleopods during coalescence, have a CR > 0.9, with CR_P2_ = 0.95 ± 0.05, CR_P3_ = 0.92 ± 0.04, and CR_P4_ = 0.93 ± 0.03 (n = 6 shrimp, Fig. 5C). The majority of the recovery stroke is dominated by pleopod coalescence. More importantly, each of these pleopods is confined between two adjacent legs for 43.6 ± 4.7, 34.8 ± 4.8, and 28.6 ± 4.2% of their recovery stroke, respectively, thus guarding individual legs from significant interactions with the surrounding fluid for about 1/3 of the recovery stroke. P1 and P5, which lead and trail the coalescing group, respectively since they only have one neighboring appendage, still contribute to coalescence for about 2/3 of their recovery stroke, given CR_P1_ = 0.76 ± 0.08 and CR_P5_ = 0.60 ± 0.06.*P. vulgaris* displays more than one metachronal wave at a given time, therefore several coalescence cycles occur within the span of a complete metachronal wave propagating from P5 to P1 (Fig. 6D). This is because the anterior legs finish their coalescence cycles while the posterior legs initiate a new metachronal wave that begins a new coalescence cycle almost immediately after. This further enhances the overall influence of pleopod coalescence during swimming.

Our robotics data emphasized how inter-pleopod phase and coalescence promotes the most consistent rearward flux and fluid momentum to maximize thrust (Fig. 7, 8, Fig. S6). When this arrangement breaks down, such as when artificially reversing the metachronal wave to induce pleopod divergence, the wake becomes more turbulent, the thrust less consistent, and individual pleopod wakes are discernable (Fig. 7, Fig. S6). To investigate how coalescence influences force production, particularly during the phase when the pleopods are closely grouped during their recovery stroke, we tested a three-legged configuration (P2-P3-P4, Fig. 8) that replicated the conditions within a coalescing group. We compared the instantaneous forces produced during the coalescence phase by the same, but independent, legs, time-synchronized to the corresponding metachronal wave of three-legged case. We found that three coalescing pleopods generate 30.2% more net thrust than independent pleopods during the corresponding coalescence phase (Fig. 8C).

**Figure 8.**
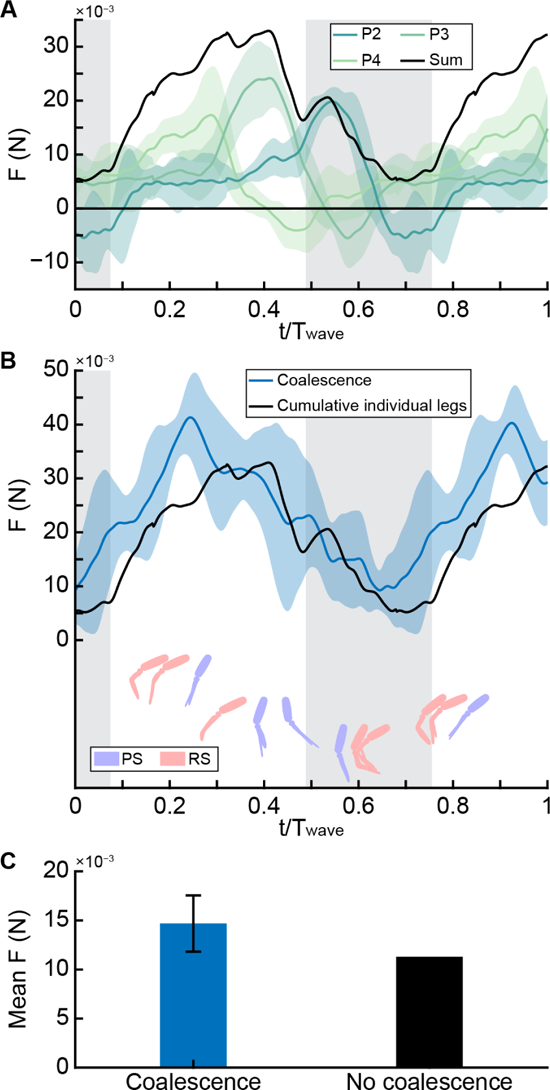
Comparing the net thrust produced by the flexible coalescing pleopod group P4-P3-P2 and the cumulative forces from the same but individual pleopods. **(A)** Instantaneous and cumulative (black) forces produced by independent legs, time-synchronized to the corresponding metachronal wave. Colored shading indicates the standard deviation for at least 8 consecutive beat cycles. Background grey shading indicates the coalescence phase. **(B)** Instantaneous thrust produced by the P4-P3-P2 coalescing group. Coalescence promotes greater net thrust during the coalescing phase (grey shaded background). **(C)** Coalescing pleopods generate 30.2% more thrust than the same but independent actuating pleopods during the coalescence phase.

## DISCUSSION

The sequential beating of coordinated appendages makes metachronal swimming an efficient mode of locomotion by producing steadier fluid flows than a single appendage to achieve the highest average flux and fluid momentum, thereby maximizing swimming performance. Several biological adaptations such as appendage spatial asymmetry and phase, and beneficial inter-appendage interactions have been identified as the main mechanisms of efficient thrust production. We found that passive bending of the swimming appendages and coalescence through metachronal coordination are also crucial in reducing drag; a determinant factor of performance in self-propelled systems. Using biological and bioinspired robotics data, we provide a comprehensive overview of the underlying mechanisms of pleopod asymmetrical bending and coalescence to explain their role in reducing drag during forward swimming.

### The role of pleopod passive bending

Appendage bending is a ubiquitous functional feature of the recovery stroke in many metachronal swimmers like copepods, ctenophores, krill, and polychaete worms. This contrasts with the power stroke during which the appendages remain relatively stiff to increase the surface area and maximize thrust. Rowing appendages are often designed, either with an articulation or with asymmetric rigidity, such that they flex posteriorly (i.e., during the recovery stroke) but not anteriorly (i.e., power stroke) (Hessler, 1985; Johnson and Tarling, 2008). In *P. vulgaris*, similar to krill, the ramal joint modulating the β angle between the rami and the protopodite can partially contribute to spatial asymmetry between the power and recovery strokes (Murphy et al., 2011; Santos et al., 2023). However, we found that the spatial asymmetry of the pleopods is primarily driven by the differential flexural stiffness along the posterior and anterior directions (Fig. 3). The rami are, on average, 1.7 times more flexible in the posterior direction — corresponding to the recovery stroke when the resistive force of the flow acts upon the anterior face.

What is the source of the observed asymmetry? Studies on ciliated microorganisms indicate that asymmetric waveforms are produced by actively sliding adjacent microtubules within the propulsor (Dutcher, 2019). This, however, is not applicable to metachronal crustaceans whose biramous pleopod rami are monolithic structures (Fig. 2, Fig. S1). To date, there is no morphological evidence suggesting the presence of structures (i.e., muscles, microtubules) capable of inducing controlled spanwise bending of the rami via active processes. Contrary to organisms living in the viscous regime, larger organisms can tune their structural (i.e., geometry) and material properties to the inertial forces they experience to achieve passive deformation under strain (Koehl, 1984; Vogel, 2020; Wainwright and Koehl, 1976). Doing so eliminates the need for active control, thus greatly simplifying actuation. Detailed μ-CT scans of the pleopods revealed that the posterior face was concave, with normalized chordwise curvature *κ* = 1.26±0.21 and 1.11±0.40 for the endopodites and exopodites respectively (Fig. 2). In plants, such chordwise geometry enables different compliances under bending with opposite orientations, with the structure bending toward the concave face (Guzmán et al., 2016; Wei et al., 2022), analogous to the recovery stroke in *P. vulgaris*. This mechanism is reminiscent of construction site measuring tapes whose chordwise curvature — which coincidentally have similar values to *P. vulgaris* — gives them strength when measuring with the concave surface facing upward and resisting gravity but high flexibility for winding in the casing. The curved geometry of the pleopod thus emerges as a passive strategy to effectively achieve spatial asymmetry. The pleopods can bend during the recovery stroke when the hydrodynamic forces resisting the motion overcome the inherently weak stiffness given by its concave shape. In contrast, this geometry stiffens the pleopods during the power stroke when resistive forces act in the opposite direction. These effects are further illustrated by the pronounced change in local spanwise curvature with pleopod angular velocity we observed only during the recovery stroke (Fig. 4). For Re >1000, drag scales with the square of the velocity of the appendages (Batchelor, 2000), making the strain due to hydrodynamic loading (i.e., bending) inherently linked with the motion of the pleopod during a beat. As such, by passively bending under fluid loading, the flexible pleopods can autonomously reduce their projected surface area to the flow and become more streamlined during the recovery stroke (Vogel, 2020).

Self-reconfiguration reduces the stress on the pleopod proportionally to the resistive fluid forces (Alben et al., 2002; Gosselin et al., 2010). We found that bending lowers the drag coefficient of individual pleopods by 41.0 to 75.8% depending on the pleopod, compared to their stiff counterpart (Fig. 5). The source of this substantial drop in the induced drag is twofold: 1) bending reduces the effective surface area to the flow, and 2) bending mitigates fluid-structure interactions, notably with the rearward-traveling power stroke wake. Because the pleopods work at Re >1000 in *P. vulgaris*, presenting a smaller profile in the flow lowers drag-inducing inertial effects and their nearly horizontal orientation further trades inertial effects for much weaker shear forces (Gosselin et al., 2010). Drag reduction is enhanced by limiting interactions with the power stroke wake that circulates nearly unimpeded posteriorly during the recovery stroke (Fig. 5A–D). For comparison, stiff pleopods systematically disrupt the power stroke wake whose kinetic energy induces substantially higher drag (Fig. 5E). Differential flexibility does not affect performance during the power stroke, as we found no difference in thrust production between the flexible and stiff treatments. The overall instantaneous force patterns during a beat are consistent with data obtained with our previous krill analog (Santos et al., 2023), but the contrast between the power and recovery strokes is stronger because we accounted for pleopod asymmetrical bending. By maximizing the thrust to drag ratio via spatial asymmetry between the power and recovery phases, shrimp can effectively overcome the resisting appendage and body drag to achieve propulsion (Byron et al., 2021; Herrera-Amaya and Byron, 2023; Lou et al., 2022; Vogel, 2020). Producing less overall drag during the recovery phase reduces the power needed to actuate the pleopod, thus promoting economical swimming and potentially lowering the cost of transport (COT) (Lionetti et al., 2023).

### The role of pleopod coalescence

The phase-dependent grouping of several closely spaced adjacent appendages — coalescence — is intrinsic to antiplectic metachronal waves (Alben et al., 2010; Byron et al., 2021; Ford and Santhanakrishnan, 2021). We found that the mean inter-pleopod phase φ = 0.18 ± 0.02 t/T and pleopod *B/L* = 0.36 ± 0.01 of *P. vulgaris* promotes the tight grouping of three leg pairs during their recovery phase and the complete separation of the legs during their power stroke (Figs. 6, 7). Many metachronal organisms display strikingly similar parameters and kinematics that fall within relatively narrow ranges — 0.2 < B/L < 0.65 and 20 < φ < 25% (Byron et al., 2021; Garayev and Murphy, 2021; Murphy et al., 2011). These commonalities are often explained in the context of the power stroke for which there is a need to synchronize the swimming appendages and circulate the wake to promote efficient force production (Ford and Santhanakrishnan, 2021; Ford et al., 2019; Murphy et al., 2013). Our robotics data support this idea that inter-pleopod phase and configuration modulates rearward average flux and fluid momentum to maximize thrust (Fig. 7, Fig. S6). When artificially reversing the metachronal wave to induce pleopod divergence during the recovery stroke, the wake becomes more turbulent, the thrust less consistent, and individual pleopod wakes are discernable (Fig. 7, Fig. S6). A recent numerical study with ctenophores found that appendage tip vortex interactions between recovering adjacent appendages are equally as important in reducing drag and can improve the thrust-to-power ratio by up to 55% (Lionetti et al., 2023). As such, by promoting inter-pleopod interactions, coalescence can have a significant impact on the performance of metachronal propulsion. This potential tradeoff between pleopod separation and grouping can explain why metachronal appendages beat within such narrow phase and *B/L* ranges.

The importance of coalescence in metachronal propulsion is highlighted by its overall pervasiveness during the recovery stroke. Coalescence was observed for over 90% of the recovery phase of the legs (Fig. 6). Note, that while all the legs display coalescence, their position and role within a coalescing group change sequentially from leading, to being confined, to trailing the group as the metachronal wave propagates. Consequently, the pleopods are exposed to changing flow conditions during their beat, which coalescence leverages to reduce the overall drag.

Pleopods P2–P4 (which have anterior and posterior neighbors) are completely confined within a group and not directly subject to the flow for about one third of their recovery stroke (Figs. 6AB, 7). To investigate how this affects force production, we tested a three-legged configuration (P2-P3-P4, Fig. 8) that replicated the conditions within a natural coalescing group. This configuration eliminates the contribution from the legs external to the coalescing group, thus strictly isolating the effects due to coalescence. Contrary to our expectations that net drag would be produced during the recovery phase of the pleopods, we found that instead, only net thrust is generated throughout the entire metachronal wave. We compared these results with the time-synchronized force data from the same three independently tested pleopods and found that even without coalescence, only net thrust is produced during a metachronal wave and fluctuates similarly over time (Fig. 8).

The inter-pleopod phase and spatial asymmetry are sufficient to ensure net thrust is always produced (Ford and Santhanakrishnan, 2021), albeit not steadily. We must inform the reader that because the robot was tethered, the results potentially slightly overestimate the forces, particularly thrust. Tethering affects the flow by artificially reducing the system’s speed to zero (Murphy et al., 2013), thus causing water to stagnate posteriorly at the end of the pleopod recovery stroke, thus partially enhancing the reactive forces of the flow in the thrust direction (swimming direction) (Santos et al., 2023). Nonetheless, coalescing pleopods produce 30.2% more mean thrust during the coalescence phase than their independent counterparts. Given that the main difference between these two cases is the strong inter-pleopod interactions only during the recovery phase of the pleopods, coalescence emerges as an effective strategy to mitigate drag and improve the overall performance. This is due in part to the strong contact between the pleopods’ surfaces forming a tight seal, shielding the coalescing group from flow instabilities that would otherwise develop between the legs (Fig. 7, Fig. S5). We observed that the recovery stroke of P5 (initiating the metachronal wave) does not shed a wake in the free stream when coalescence occurs. Instead, the posterior bound vortex is consistently circulated behind the next anterior pleopod as the coalescence group propagates to P1. The traveling recovery stroke bound vortex was then trapped between the pleopods and the much stronger inferior, rearward-traveling power stroke wake into which it dissipates at the end of the metachronal wave (Fig 7). This mechanism is consistent with the vortex-weakening mechanism found in ctenophores that harnesses fluid-structure interactions among closely spaced appendages to reduce drag (Lionetti et al., 2023).

Overlapping the profile of several appendages via coalescence can reduce the cumulative effective surface area of multiple appendages arranged in series. But does coalescence turn the drag of three otherwise separate pleopods to that of only one? We found that coalescence does not induce net drag at all, unlike individual legs that all produced net drag during their recovery phase (Figs. 5, 8). This results from the combined effects of the phase and resulting coalescence. The inter-pleopod phase intrinsically synchronizes the forces produced by the pleopods such that net thrust is always generated. However, coalescence cannot exist without the appropriate phase, and we could not strictly isolate the independent contribution of pleopod grouping (i.e., overlapping profiles) and the three phases of coalescence to lowering the drag. Future experiments designed to measure the instantaneous force output by each leg within a group could help clarify how coalescence affects their performance relative to their independent counterparts.

### Pleopod bending and coalescence are complementary

Pleopod coalescence is intrinsically linked with the *B/L* ratio, the inter-pleopod phase, and beat amplitude. Coalescence occurs by virtue of the spatial and temporal synchronization of several appendages when two or more adjacent appendages are oriented in opposite directions (i.e., posteriorly during the power stroke and anteriorly during the recovery stroke). Thus, coalescence can occur even in the absence of appendage bending. However, flexible pleopods leverage coalescence by enhancing structure-structure and fluid-structure interactions. In addition to the mechanical and hydrodynamic benefits of bending, they are not as limited by the available space between them and physical interactions as their stiff counterparts, thus enabling wider stroke amplitude (Brennen, 1974). This gain in the plasticity of the motions of the pleopods means that physical interactions are not detrimental and might instead be instrumental in promoting constructive physical interactions, such as when the legs coalesce.

While stiff rami can still coalesce, they achieve only partial spanwise contact, often resulting in strong physical interactions that disrupt the geometry of other pleopods during the recovery stroke. These undesired effects induce a turbulent wake and cause significant disturbances in the instantaneous forces produced during coalescence (Fig.7, S6C). In contrast, spanwise bending enables the precise overlapping of several pleopods such that they conform to each others shape and form a tight seal within a coalescing group to prevent inter-pleopod flow (Fig. 6,7, S5). This reduces the impact of the recovery stroke on the overall wake and promotes more consistent thrust production (Fig. S6A). One potential challenge posed by the close contact of adjacent pleopod is the need to overcome capillary adhesion when separating the trailing pleopod from a coalescing group. Because of surface tension, a thin layer of water is confined between two adjacent legs, acting at the interface between the fluid and the wetted surface. At intermediate Re, the capillary effects can be significant (Butler and Vella, 2020; Lambert, 2013), potentially rendering coalescence less effective.at the moment of pleopod separation. We found that having flexible pleopods mitigates this problem. The trailing and confined pleopods in a coalescing group (see Fig. 6 for reference) slide against one another and the rami of the trailing leg unfurl progressively along their span during the later phase of the recovery stroke due to direct physical interaction with the anterior leg (Fig. 6, S5). This mechanism is particularly noticeable in Fig. 4A where the instantaneous local spanwise curvature maximum travels from the proximal to the distal section of the rami and is greatest near the tip during the late phase of the recovery stroke. Unlike two rigid flat plates that induce the highest capillary effect when pulled apart in opposite directions, unfurling the trailing pleopod localizes the capillary force at the point of separation rather than the entire surface (Butler and Vella, 2020; Lambert, 2013), thus reducing the force required to separate from the coalescing group. Pleopod bending is crucial to enhance structure-structure and fluid-structure interactions and leverages coalescence from a simple byproduct of inter-pleopod phase, separation and amplitude, to a mechanism optimized for minimizing disruptive hydrodynamic effects, thereby improving overall swimming performance.

While *P. vulgaris* displays perfectly overlapping pleopods during coalescence, appendage contact is not always present in other organisms like ctenophores (Colin et al., 2020; Herrera-Amaya and Byron, 2023) and ciliates (Blake, 2001; Blake and Sleigh, 1974; Sleigh, 1989) whose appendage Re is substantially lower (Re < 100). Still, it is interesting that all the metachronal swimmers studied to date display some form of flexible appendages that can bend during their recovery stroke, leading to the formation of an array of somewhat overlapping appendages. At higher Re, this configuration combines the reduced total effective surface area of individual appendages to the flow due to bending and the masking effect from the anterior appendage to reduce drag (Alben et al., 2002; Barsu et al., 2016; Gosselin et al., 2010). By enabling close physical contact between the pleopods, shrimp mitigate parasitic flow features that would develop within a loose pleopod cluster. At low Re where viscous forces become more important, the need to prevent inter-appendage flow is not as crucial given these instabilities dissipate quickly. However, achieving appendage overlap during their recovery stroke by maintaining adequate appendage spacing, phase, and amplitude, may serve in reducing surface drag (i.e., friction) and can promote fluid interactions that can reduce drag and improve the thrust-to-power ratio (Lionetti et al., 2023).

## CONCLUSIONS

Metachronal swimming is commonly found across many taxa and a wide range of scales, suggesting common biophysical mechanisms driving its success. The performance of metachronal propulsion is generally perceived from the need to maximize thrust production. However, we found that asymmetrically flexible appendages and coalescence (major commonalities among these organisms) also enhance the overall swimming performance by significantly reducing drag. Using experimental methods, we present a comprehensive overview of the underlying morphological, functional, and physical mechanisms to quantify the drag-reduction capabilities of pleopod flexibility and coalescence. We found that the curved cross-sectional profile of the pleopods enabled passive asymmetrical bending during the recovery stroke to reduce the coefficient of drag of each pleopod by 41 to 75.8% relative to stiff pleopods. In the context of metachronal swimming when up to three pleopod pairs interact, coalescence reduces the overall drag of the group such that the mean net thrust produced during the coalescence phase is increased by 30.2%. Nonetheless, because we could not resolve the forces for each pleopod within a coalescing cluster during the transition from leading, to confinement, to trailing the group, the question whether the drag of three overlapping pleopods becomes that of only one remains open. Future robotics experiments, including untethered swimming, might provide the answer and help quantify the energetic benefit of pleopod bending and coalescence to demonstrate their role in achieving high swimming performance and efficiency. Our current result improves our understanding of the fundamental biological and physical components of metachronal propulsion that may aid the development of novel bio-inspired underwater metachronal vehicles.

## Acknowledgements

We are grateful to Erika Tavares for performing the μ-CT scans of *P. vulgaris*. We also gratefully acknowledge Adrian Herrera-Amaya for his insightful discussions that strengthened the foundation of this work. We thank Marjorie Bradley for her assistance with coordination and administrative support throughout the duration of this research. Funding for this research was provided by the National Aeronautics and Space Administration (NASA) Rhode Island Established Program to Stimulate Competitive Research (EPSCoR) Seed Grant.

