## Supplemental methods and figures for "Going around the bend to understand the role of leg coalescence in metachronal swimming"

### 1. μ -CT scan and three-dimensional reconstruction

A SkyScan 1276 high-resolution microtomography (Bruker, Billerica, MA, USA), upgraded to receive a Hamamatsu microfocus X-ray source (L10321-67, Bridgewater Township, NJ, USA) and a 4K camera (XIMEA MH110XC, Lakewood, CO, USA) was used to perform high-resolution scans. The scanning parameters were set up as follows: Isotropic voxel size = 10.287245 μm per pixel; Source voltage = 45kV, Source current = 200μA; Rotation step = 0.2º. We used the Bruker micro-CT Skyscan software (NRecon, DataViewer, CTAnalyser, Version 2.0.0.5) for primary 3D reconstructions from cross-section images. Volume rendering images were obtained with Slicer v.5.0.3. It is important to point out that Slicer's volume renderings were obtained by loading mirrored raw images obtained with the Bruker software because the latter outputs mirrored, inverted images, flipping the right and left sides of the specimen.

### 2. Measuring *P. vulgaris* pleopod chordwise curvature

The 3D reconstructions of the μ-CT scan of ten pleopods of a shrimp specimen were used to obtain mid-length chordwise transections of each endopodite and exopodite (Fig. S1A,B). We measured chordwise curvature $R$ by applying the Menger curvature of a triple of points, which we defined as two points located at the extremities of the curved chord (from cross sections in Fig. S1B) and a third point located along the chord.

$$R = \frac{abc}{4S} \quad (1)$$

where $a$ and $b$ are the segments joining the extremities to the center point along the chord, $c$ is the segment formed by the two extremities (see Fig. S1C), and $S$ is the surface area of the triangle. Chordwise curvature was normalized using the maximum effective width $W$ of the appendage (i.e., the distance between the two points at the extremities of the chord) as $\kappa = W/R$.

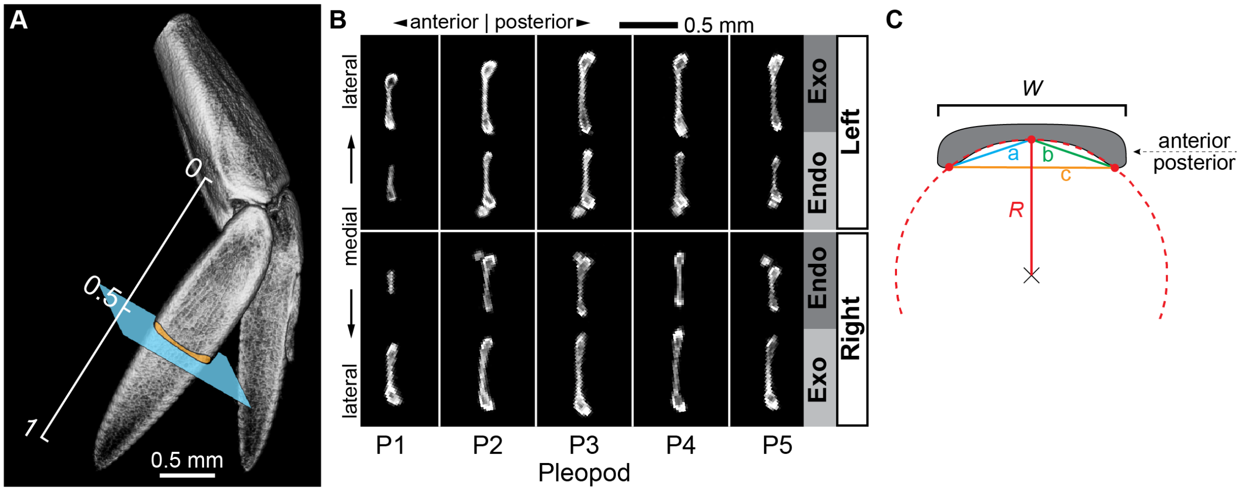


**Figure S1: Pleopod cross sections from μ-CT scans showing the curved chordwise profile of the ramal structures. (A)** Cross sections were taken at the mid-length (normalized length = 0.5) of the endopodites and exopodites of the ten pleopods of one *P. vulgaris* specimen. **(B)** The posterior face of each ramus had a pronounced chordwise curvature. Note the presence of the appendix interna along the median edge of the endopodite (showing as a small circle in cross sections). This small structure is used by shrimp to temporarily connect the left and right endopodite during swimming. **(C)** Measuring the chordwise curvature of the rami from three points located along the posterior face. *W* is the width of the ramus and a, b, and c are the segments between three points selected along the posterior profile to calculate the radius of curvature *R* of the fitted circle.

### 3. Calculating the depth of focus

The depth of focus (DoF) of the present in-vivo bright-field PIV system was evaluated according to (Nakano et al., 2005):

$$DoF = \frac{n\lambda}{NA^2} + \frac{ne}{M\,NA} \tag{2}$$

where *n* is the refractive index of the fluid medium (≈ 1.34 for our experiments), $\lambda$ is the source light wavelength (≈ 0.53 μm), *NA* is the numerical aperture of the 60 mm macro lens (*NA* = 1/5.6 = 0.18), *M* is the magnification of the true lens objective (Nikon specify that this lens has a 1:1 magnification), and e is the camera sensor pixel size (i.e., the smallest resolvable distance = 6.5 μm). The calculated DoF is ≈ 57.1 μm.

### 4. Five-legged robotic analog of *P. vulgaris*

We designed the 20× scaled five-legged robot as a modified version of the Pleobot developed by (Santos et al., 2023). The architecture and operation of each leg is similar to that of the Pleobot. Each leg is controlled by two servo motors: one controlling the α angle and the other driving the β angle (Fig. S2A). Because of this design, the β angle is not strictly independent from the α actuation. Indeed, without correcting for the α motion, the β angle would actuate with the same magnitude but in the opposite direction to the α angle. Thus, we computed the effective β kinematics by subtracting the effects of the α displacement. The main difference in our version of the robot is the use of pulleys rather than a gear train. This has the advantage of preventing backlash, reducing the part count, and lowering the noise otherwise caused by imperfections in the 3D-printed gears. Therefore, the pulley system improved the accuracy and reliability of the leg actuation to achieve the desired kinematics consistently (Fig. S2B). The diameter of the driving pulleys (mounted on the servos) was twice that of the driven pulleys (within the protopodite, at the α and β joints), giving a gear ratio of 0.5 (amplification). The accuracy of pleopod actuation was about 2°, which is consistent with the precision of the servomotors (accuracy ≈ 1°, HS-81, Hitec, USA) and the pulley ratio. Another benefit of the pulleys is the ability to modify the length of the protopodites to match *P. vulgaris* morphometrics by simply changing the distance between the pulleys.

Each servo motor was calibrated by visually mapping the range of travel because the effective range of motion can differ substantially from the factory specifications provided by the manufacturer. Pictures of the servo motors in their minimum and maximum positions were analyzed in ImageJ to measure the total range of motion (error = 0.1°). This step was necessary to ensure that the kinematics data fed to the servo (as a pulse width, in ms) corresponded perfectly with the desired rotation of the servomotor. For instance, considering the pulley ratio of 0.5, every 1° kinematics step (and, by extension, the motion of the actual pleopod) had to result in a 0.5° rotation of the servomotor.

After the servo motors were mounted on each leg, they underwent a second phase of calibration. This is because the median position of the servo may not precisely match the neutral position of the pleopod — considered here as fully extended vertically. Each servo was assigned a correction factor to zero its median position. The final kinematics data input to the servomotors of the five legs, based on the kinematics of *P. vulgaris*, was calculated by accounting for the effective servo range and correction factor. The motion program was then fed to an Arduino microcontroller (Mega 2560 Rev3, Arduino, Italy). While we calculated the mean α kinematics for six specimens, we used the kinematics data measured for only one representative forward steady swimming shrimp (cruising horizontally) to drive the robotic pleopods motions. We did this because the timing and position of the pleopods relative to one another are key to metachronally beating closely spaced legs without destructive physical interference. Averaging the α angle over time for several individuals masks the subtle differences in leg motions (temporally and spatially) between individuals due to some variability in the swimming speed, total stroke amplitude, and pleopod phase. Using such averages caused undesired physical contact and interferences between the protopodites. As such, we used the kinematics data for one representative shrimp, which displayed horizontal, forward steady swimming. Two complete consecutive beat cycles were extracted from the high-speed video and averaged to smooth out the measurement uncertainty during manual tracking.

We fitted the pleopods with flexible ramal structures (Practi-Shim™, thickness = 0.0254 mm), whose chord profile was heat-shaped using the same curvature profile identified from μ-CT scans of shrimp pleopods. This thickness was selected because, when combined with the corresponding chordwise curvature, it resulted in qualitatively comparable spatial asymmetry to *P. vulgaris* rami.

Heat shaping was performed by pre-heating a small rectangular plastic shim using a heat gun. This was then clamped to cool down in a custom-made 3D-printed mold whose dimensions were specific to the specified chordwise curvature (Fig. S2C,D). The final shape of the rami was cut using stencils whose widths were adjusted for the chordwise curvature of each ramus to produce the correct effective ramus width. We applied a ramus width correction factor ($C_E$) to the stencil to maintain the natural pleopod effective surface area, accounting for the effective width reduction caused by curving the chord. Because the profiles of the endopodite and exopodite were established using the top-down view of a *P. vulgaris* pleopod, we only captured the effective width of the structures rather than the length of the curved chord (arc length). Using the effective width rather than the arc length inevitably yields incorrect chordwise dimensions upon shaping the ramal structures. To account for the added width when designing the stencils, we expanded the profiles laterally using the maximum effective width of the endopodites and exopodites ($W$). The expansion coefficient was calculated as follows:

$$C_E = \frac{2\pi R\left(\left(2sin^{-1}\left(\frac{W}{2R}\right)\right)/360\right)}{W} \quad (3)$$

Where $R$ is the radius of curvature (the radius of the heat-shaping mold), and $W$ is the maximum effective width of the pleopod. The proximal section of each ramus was finally glued to the respective ramus stem (endopodite or exopodite) using cyanoacrylate glue (Fig. S2A). The stiff rami were made following the same heat-shaping method and chordwise curvature but with thicker plastic shims (Practi-Shim™, thickness = 0.1016 mm). The stiff rami were still flexible enough to prevent destructive interactions between legs that would damage the robot when the legs collided but remained completely stiff during the pleopod recovery stroke and when mild pleopod contact occurred. These rami did not bend significantly during coalescence either.

We recognize that there are several limitations to the current design. While we reliably achieved pleopod bending and the passive actuation of the γ angle to change the surface area of the pleopods during a beat, we could not include the setae. In *P. vulgaris*, the setae spread during the power stroke to substantially increase the surface area of the membranous part of the rami. In contrast, they fold completely along the leading and trailing edges of the rami during the recovery stroke. While our results may underestimate the thrust coefficient during the power stroke, we focused exclusively on pleopod bending and coalescence during the recovery stroke in this investigation. In this context, by only including the membranous part of the rami, we achieved analogous functional morphology as *P. vulgaris* during the recovery stroke. Also, note that the robot was tethered to the force transducer to measure the thrust and drag forces. While the literature reports that tethering can affect the flow fields by restraining the robotic model, we still reliably capture the principal flow patterns produced by *P. vulgaris* and estimate the forces produced by and acting upon the pleopods.

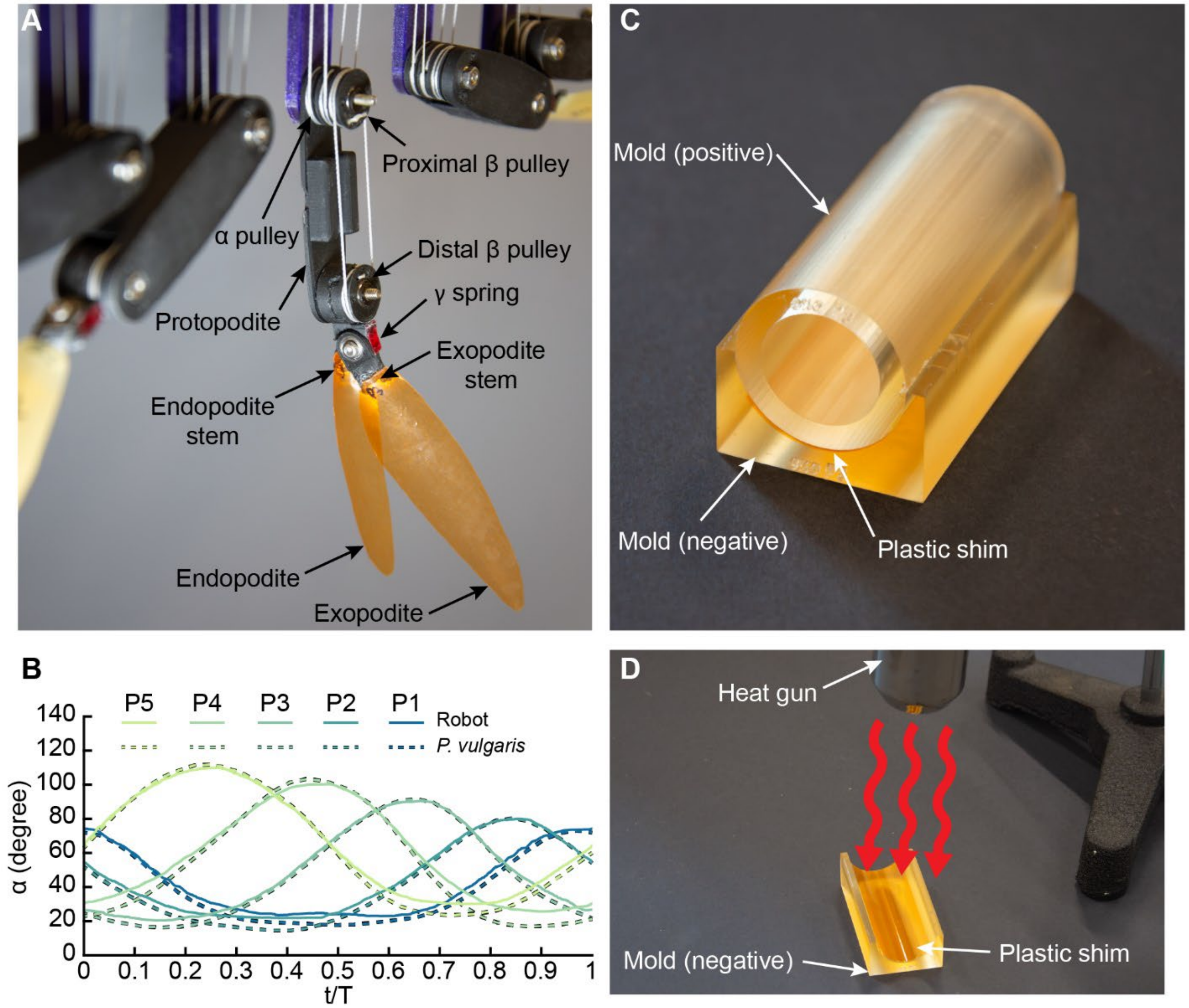


**Figure S2. Design, fabrication, and kinematics of the five-legged robot analog. (A)** Architecture of the robotic legs (without protopodite cover). Two separate servomotors control the α and β angles. α is controlled by the α pulley mounted in the proximal joint of the protopodite. β actuation is achieved with two pulleys. The servomotor is linked to the proximal β pulley (mounted on the same axle as the α pulley), which is linked to the distal β pulley. This design allows the independent actuation of the α and β angles. **(B)** Comparison of robotics and biological pleopod kinematics. The motions of the robotic legs (averaged over three consecutive strokes) were driven by the mean α amplitude of the five pleopods of one representative steadily swimming *P. vulgaris* cruising horizontally (obtained from two consecutive beats). Note that the standard deviation for the robotics data (shading) blends with the curves for the mean data because the actuation accuracy is within ≤ 2°. **(C)** The curved rami were produced by shaping the plastic shim with the desired chordwise curvature using positive and negative molds (here, for the P3 exopodite). **(D)** Heat shaping the plastic shim making up the flexible rami (here, for the P3 exopodite).

### 5. Matching the beat frequency of the robot to Re = 1720

The robot was dynamically and kinematically scaled to achieve the same flow regime as *P. vulgaris* during forward steady swimming (matching Re ≈ 1720 achieved by *P. vulgaris*, measured using pleopod P3). First, we dynamically scaled the experiment using a 3:2 glycerin-water mix ($\rho = 1160$ kg m$^{-3}$, $\mu = 0.0114$ kg m$^{-1}$ s$^{-1}$, $\nu = 9.8652\times10^{-6}$ m$^2$ s$^{-1}$). This is because using water alone would require the kinematics to be adjusted to match *P. vulgaris* Re such that the beat frequency would fall well below what the servomotor can accurately output (see (Santos et al., 2023). We estimated that this glycerin mix would enable beat frequencies $f = 0.1$ to 2, well within the specifications for the servos and experimentally relevant for high-speed PIV. To further match the natural Re = 1720, we performed kinematics adjustments, particularly the beat frequency (dictating the pleopod tip velocity). To achieve the desired pleopod beat frequency and tip velocity needed to reach a Re = 1720 in the 3:2 glycerin-water mix, the number of servo steps was calculated as follows:

For $\nu_{glycerin/water} = 9.8652\times10^{-6}$ m$^2$ s$^{-1}$, the expected pleopod maximum tip velocity, $U_{max} = 0.16757$ m s$^{-1}$ per equation (4):

$$U_{max} = \frac{9.8652\times10^{-6} Re}{L} \tag{4}$$

where L is the 20× scaled robotic protopodite + exopodite P3, in m. The beat frequency could only be adjusted by increasing the number of kinematics steps the microcontroller outputs to the servos within a full beat cycle, because the time between steps was fixed to 20 ms for the particular servos we used (HS-81, Hitec, USA). The number of steps driving the servos during a beat cycle dictates $U_{max}$ of the pleopod according to the power function: $U_{max} = 10.688\, steps^{-0.85}$ ($R^2 = 0.974$). This was obtained by measuring the kinematics of the robotic pleopod for nine different servo step counts ranging from 10 to 80 points using high-speed video (Fig S3A) and fitting a power function to the data using Matlab 2023b. Using this relationship, we determined the ideal servo step number $n_i = 133$ to achieve Re = 1720 from the following equation:

$$n_i = \sqrt[0.850]{\frac{10.688}{U_{max}}} \tag{5}$$

From the same videos, we measured the beat frequency as $f = 1/t_{stroke}$, where $t_{stroke}$ is the total time to complete a full beat. Using $n_i$, we found the theoretical beat frequency of the legs $f = 0.374$ s$^{-1}$ (Fig. S3B). We measured the actual beat frequency of the legs from the experimental high-speed PIV videos presented in this investigation as $f = 0.375$ s$^{-1}$. We used this experimental beat frequency to calculate the thrust and drag coefficients.

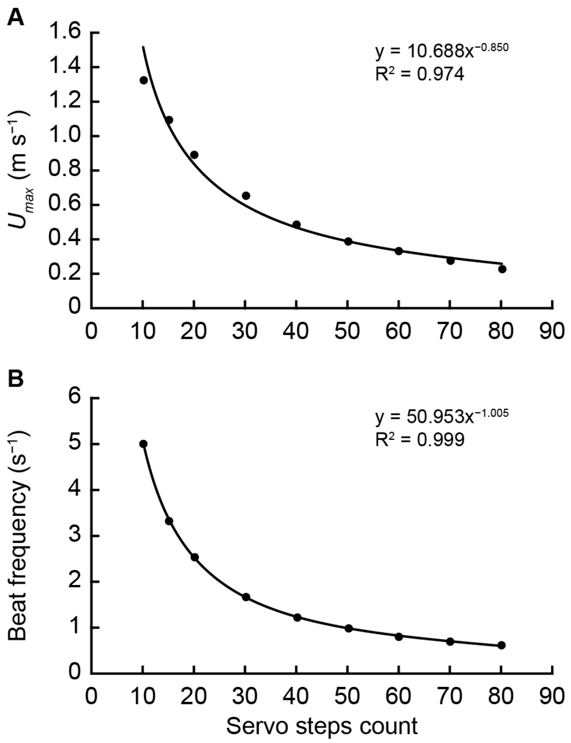


**Figure S3. Relationship between servo step count and the pleopod maximum tip velocity and beat frequency. (A)** $U_{max}$ was measured from high-speed videos to inform which step count yields the proper speed to achieve Re = 1720. **(B)** The beat frequency is dictated by the number of servo steps. To achieve Re = 1720 in the 60:40 glycerin-water mix, the servos driving the robotic pleopod needed 133 steps and achieved a beat frequency $f = 0.375\ s^{-1}$.

### 6. Statistical results — Pleopod flexural stiffness

**Table S1. Statistical results of the Mann-Whitney *U*-tests to compare the flexural stiffness of the posterior and anterior faces of the endopodite and exopodite of each pleopod.** Note that the P4 endopodite of one specimen was removed from analyses because it showed significant damage after placing it in the visualization vessel.

| Ramus | Pleopod | *P*-value | n |
|---|---|---|---|
| Exopodite | P1 | 0.008 | 5 |
| | P2 | 0.056 | 5 |
| | P3 | 0.016 | 5 |
| | P4 | 0.095 | 5 |
| | P5 | 0.032 | 5 |
| Endopodite | P1 | N/A | 0 |
| | P2 | 0.032 | 5 |
| | P3 | 0.222 | 5 |
| | P4 | 0.343 | 4 |
| | P5 | 0.032 | 5 |

### 7. Passive asymmetrical bending of all the pleopods

The instantaneous local spanwise curvature travels from the proximal to the distal region during the recovery stroke (RS) in all the pleopods (Fig. S4). No significant spanwise bending was observed during the power stroke. However, the posterior pleopods (P4 and P5) show some distal curvature at the beginning of the power cycle. This is because the rami had not fully unfurled by the time the protopodite started a new beat (see Fig. S5). The transition from PS to RS was recorded as the increase in the $\alpha$ angle after reaching its minimum at the end of the RS. This is highlighted by the blue P3 instantaneous profiles that show that P3 remained stiff during this phase. In contrast, a wave of curvature travels from the proximal to the distal section of the ramus during the recovery stroke, as seen with the substantial increase in local curvature over the length of the ramus, over time.

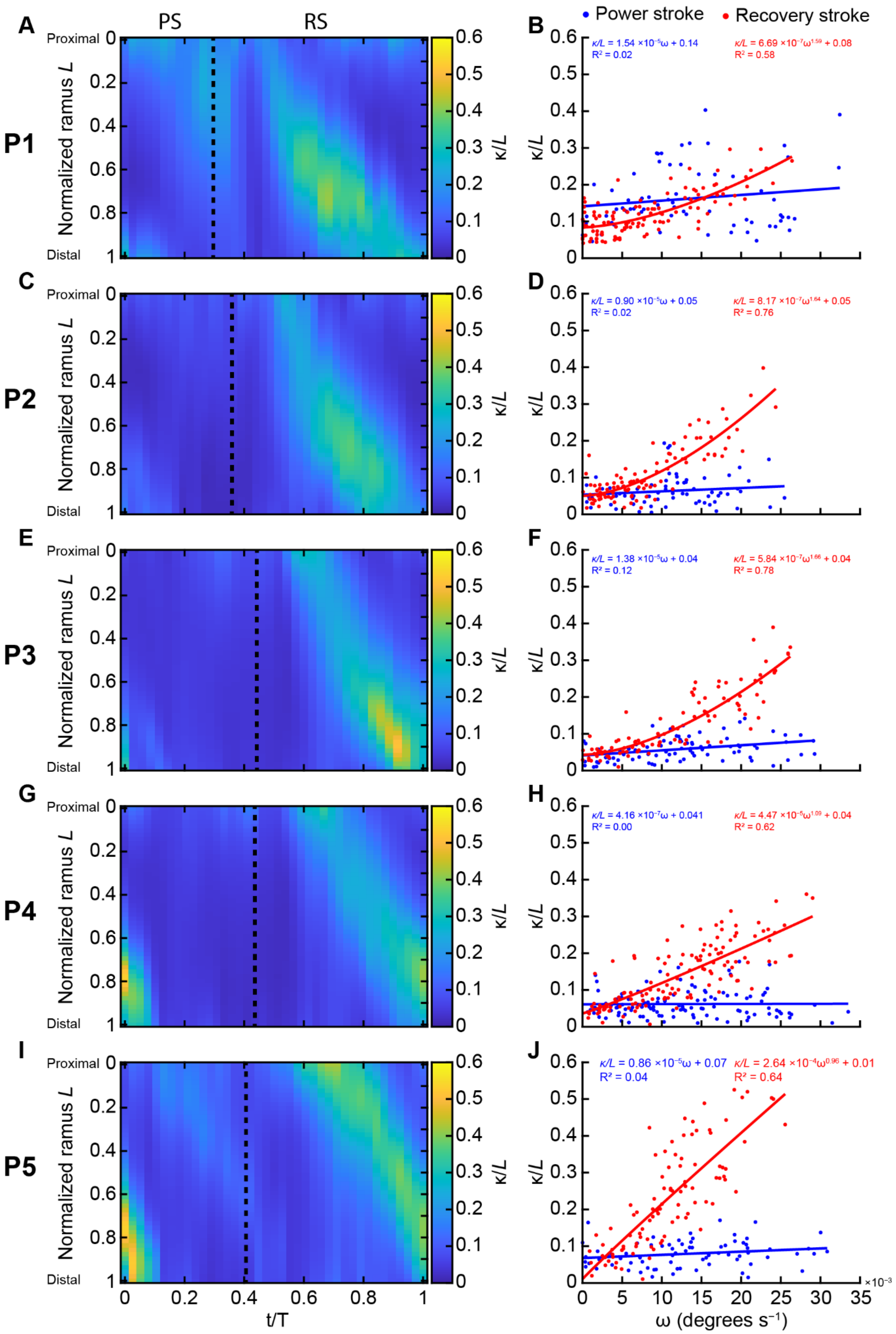

A
PS
RS
P1
Proximal
Distal
Normalized ramus L
κ/L
B
Power stroke
Recovery stroke
κ/L = 1.54 ×10⁻⁵ω + 0.14
R² = 0.02
κ/L = 6.69 ×10⁻⁷ω^1.59 + 0.08
R² = 0.58
C
P2
D
κ/L = 0.90 ×10⁻⁵ω + 0.05
R² = 0.02
κ/L = 8.17 ×10⁻⁷ω^1.64 + 0.05
R² = 0.76
E
P3
F
κ/L = 1.38 ×10⁻⁵ω + 0.04
R² = 0.12
κ/L = 5.84 ×10⁻⁷ω^1.66 + 0.04
R² = 0.78
G
P4
H
κ/L = 4.16 ×10⁻⁷ω + 0.041
R² = 0.00
κ/L = 4.47 ×10⁻⁵ω^1.09 + 0.04
R² = 0.62
I
P5
J
κ/L = 0.86 ×10⁻⁵ω + 0.07
R² = 0.04
κ/L = 2.64 ×10⁻⁴ω^0.96 + 0.01
R² = 0.64
t/T
ω (degrees s⁻¹)
×10⁻³

**Figure S4. Passive asymmetrical bending of the five pleopods (P1–P5) during a beat. (A,C,E,G,I)** Local curvature along the normalized length measured along the flexible exopodite of the corresponding pleopod during a full beat cycle. Note that curvature is normalized to the ramus length $L$. Note the progression of the instantaneous local spanwise curvature from the proximal to the distal region during the recovery stroke (RS, right of the vertical dashed line) in all the pleopods. No significant spanwise bending was observed during the power stroke. However, the posterior pleopods (P4 and P5) show some distal curvature at the beginning of the power cycle. This is because the rami had not fully unfurled by the time the protopodite started a new beat. The transition from PS to RS was recorded as the increase in the $\alpha$ angle after reaching its minimum at the end of the RS. **(B,D,F,H,J)** Local spanwise curvature at $0.3L$ of the exopodite of the corresponding pleopod as a function of angular velocity during a full beat (n = 6 shrimp). The local curvature increases proportionally with the angular velocity of the pleopod during the recovery stroke following a power relationship ($\kappa/L = a\omega^b + c$). In contrast, the rami bend only very little with increased pleopod angular velocity during the power stroke, which is best modeled using a linear fit ($\kappa/L = a\omega + b$).

### 8. Pleopod sliding and unfurling during coalescence

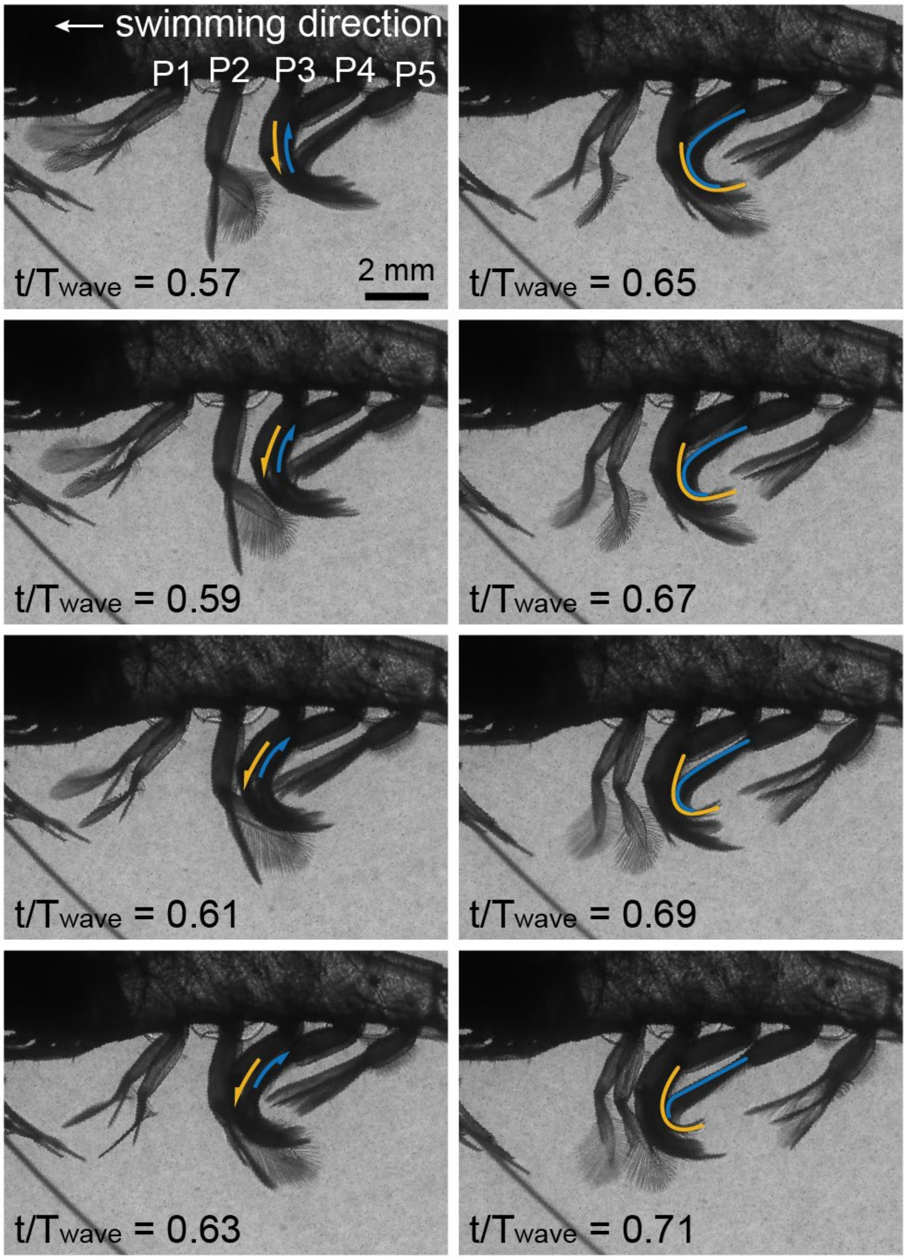


**Figure S5. Interactions between the confined and trailing pleopods during coalescence.** Initially, the trailing (orange) and confined (blue) pleopods in a coalescing group slide against one another ($t/T_{wave}$ = 0.57–0.63). Then the rami of the trailing pleopod eventually unfurl progressively along their span during the later phase of the recovery stroke due to direct physical interaction with the anterior, confine pleopod ($t/T_{wave}$ = 0.65–0.71). The trailing pleopod peels off of its anterior neighbor at the area of maximum local spanwise curvature, which travels from the proximal to the distal section of the rami over time. Time is normalized to the metachronal wave period, $T_{wave}$, defined as the time between the beginning of the power stroke of P5 (initiating the wave) and the end of the recovery stroke of P1 (ending the wave), after all the other pleopods have completed their cycle.

### 9. Comparison of the three-legged trials

While we did not emphasize the results we obtained when testing coalescence using stiff robotic appendages, there are notable undesirable resulting effects we want to point out to the reader. The qualitative comparison between the four cases involving flexible or stiff and coalescing or diverging (180° out of phase metachronal wave) pleopods highlights the benefits of flexibility during coalescence (Fig. S6A). Because physical interactions are enhanced by flexibility — it enables pleopods to wrap around one another — bending also prevents destructive physical interactions as the posterior pleopod coalesces with its anterior neighbor. The sharp oscillations observed with stiff coalescing pleopods during contact (gray background shading in Fig. S6C) result directly from the hard contact between the pleopods. Because it is rigid, the posterior pleopod (here P4) rammed into its adjacent neighbor (P3), causing instabilities in the motions (i.e., the exopodite was forced to abduct) and inducing flow instabilities (Fig. 7). While coalescing stiff pleopods still generate some net thrust, the forces are sporadic and likely undesirable to maintain efficient, steady swimming. Note that the stiff appendages we used in the robotic experiments were semi-rigid. We designed the stiff rami such that if the force were too excessive, the material could still bend partially to avoid the potential destruction of the legs. This was a preventive measure that did not influence our results. We did not observe pleopod bending with stiff coalescing pleopods, in part because contact between adjacent legs caused the rami to abduct instead.

Interestingly, when the metachronal wave was reversed and caused pleopod divergence during their recovery stroke, axial force in the swimming direction (net thrust) was maintained despite having individual pleopods interacting individually with the flow (Fig S6B). With flexible pleopods, this can be explained by the far lower drag induced when the pleopods bend, which when coupled with phase, enables more total thrust to be generated than drag. With stiff appendages, we see that little net thrust is produced during the recovery phase of the pleopods and drag dominates ($t/T_{wave} = 0.6–0.85$, Fig. S6D). This is because stiff appendages have a comparable drag coefficient during the recovery and power stroke. In this case, diverging pleopods induce maximum thrust as they interact with water individually.

Note that overall, coalescence and pleopod flexibility have only marginal effects on thrust production, even when the pleopods coalesce during their power phase when the metachronal wave is reversed. This is explained by the fact that even when the legs coalesced during the power stroke, they did not cause the same level of destructive interactions as coalescence during the recovery stroke. In fact, such interactions may promote the passive abduction of the exopodite during the power stroke, which is expected anyway during the power phase.

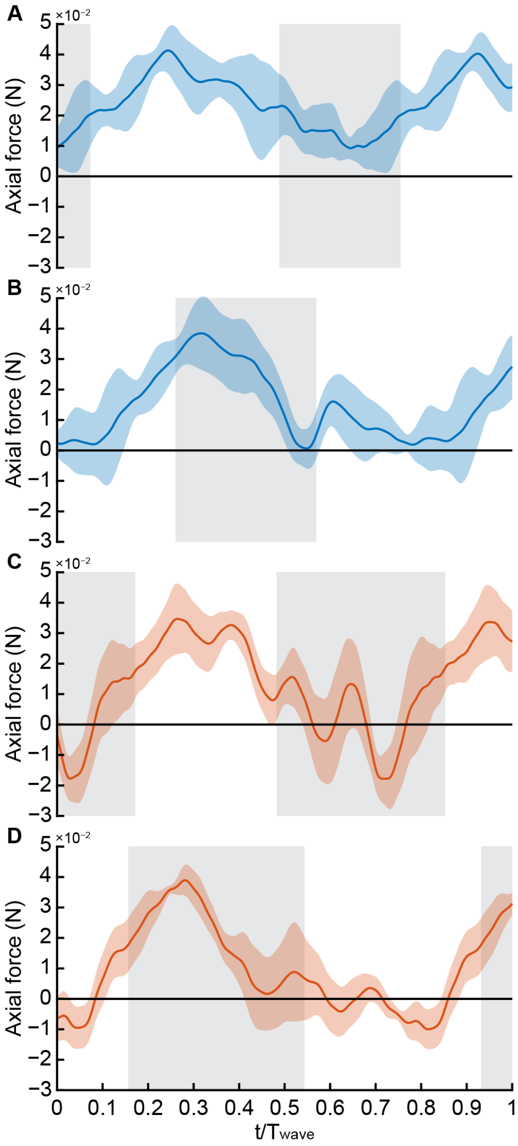


**Figure S6. Comparison of the instantaneous thrust produced by the P4-P3-P2 pleopods group over a coalescence wave with flexible or stiff ramal appendages.** Four cases are represented by **(A)** natural kinematics and flexible rami, **(B)** Reversed kinematics with flexible rami, **(C)** natural kinematics with stiff appendages, and **(D)** reversed kinematics and stiff rami. Reversed kinematics was achieved by offsetting the metachronal wave by a phase $\varphi = 180°$. Colors indicate whether the rami are flexible (blue) or stiff (orange). The data is averaged for at least 8 consecutive beat cycles. The shaded grey background indicates the time when coalescence occurs during the recovery stroke with normal kinematics **(A,C)** and contact between the legs during the power stroke with reversed kinematics **(B,D)**. Note that coalescence is defined only in the context of the recovery stroke when the pleopods wap around one another due to their close proximity. Time is normalized to the metachronal wave passing through the three pleopods making up the representative coalescing group as $t/T_{wave}$. The wave period extends to the first pleopod starting its power stroke to the last pleopod finishing its recovery stroke.
